# Maternal diet and genetics shape the human milk metabolome

**DOI:** 10.64898/2026.08.11.744248

**Authors:** Kelsey E. Johnson, Yiran Duan, Andrew Youssef, Juan J Aristizabal-Henao, Abigail Johnson, Michael A. Kiebish, Emily M. Nagel, Kristin Palmsten, Stephanie Pierce, Sarah Wernimont, Lars Bode, Eric F. Lock, Elvira M. Isganaitis, David A. Fields, Frank W. Albert, Ran Blekhman, Ellen W. Demerath

## Abstract

Human milk contains a diverse array of metabolites that contribute to infant nutrition, immune development, and microbial colonization. The maternal factors shaping the milk metabolome, and the relative contribution of genetics or diet vs. other factors, remain poorly understood. Here, we profiled 458 milk metabolites in 349 one-month postpartum human milk samples and integrated metabolomic data with maternal diet, clinical, transcriptomic, and genomic measurements. Maternal diet was broadly associated with milk metabolite composition, with significant correlations identified between dietary features and 323 metabolites. Coffee consumption strongly predicted milk quinic acid and 1,3-dimethyluric acid abundance, while high-fiber dietary patterns were associated with metabolites including proline-betaine and N-acetylornithine. Integration of milk transcriptomic and metabolomic data via machine learning identified biologically plausible gene-metabolite pairs, including associations between *QPRT* expression and quinolinic acid, and *DPEP1* and cysteine-glycine dipeptide. Genome-wide association analyses identified nine study-wide significant metabolite quantitative trait loci, including novel milk-specific associations near *PDE6A* affecting purine metabolites and near *GNE* affecting free sialic acid. Comparison with plasma metabolite studies demonstrated both shared and milk-specific genetic regulation of metabolites. Finally, we found that of all tested maternal features, diet explained the largest proportion of variation in the milk metabolome. Together, these findings demonstrate that the human milk metabolome reflects both maternal exposures and mammary gland-specific biology. This work establishes a framework for understanding how genetic and environmental factors shape milk composition.

## Introduction

Nutrition during the first 1000 days is critical to infant development and health. Human milk contains critical components that support the unique postnatal growth and development needs of the human infant, including a variety of nutritive and non-nutritive elements (e.g., fatty acids, amino acids, oligosaccharides)^1^. Milk composition varies across individuals, populations, and over the course of lactation. For example, certain fatty acids in milk are strongly influenced by maternal diet^2^. However, other components, such as lactose^3^, are tightly regulated and exhibit low variation across individuals. Maternal factors including genetics, environmental exposures, diet, and other lifestyle factors all play a role in shaping the composition of milk, but their relative contributions are largely unexplored.

The milk metabolome includes small molecules involved in energy production, amino acid metabolism, lipid metabolism, xenobiotic processing, carbohydrate metabolism, and numerous other cellular processes^4^. Maternal factors such as gestational diabetes status, BMI, diet, environmental exposures, and lactation stage have been linked to the milk metabolome^5–7^. Because metabolites represent the downstream products of gene regulation, enzyme activity, nutrient availability, and environmental exposures, the metabolome provides a high-dimensional snapshot of mammary gland physiology and thus an ideal assay to explore the respective contributions of maternal factors to milk synthesis.

The physiology of milk production is key to understanding how maternal factors shape milk composition. Some components, such as lactose and human milk oligosaccharides (HMOs), are synthesized *de novo* in the mammary gland^8^, while others are transported from circulation (e.g. micronutrients^9^), or a combination of the two (e.g. fatty acids^10^). Genetic variation that influences the milk-producing mammary epithelial cells can thus directly impact those components synthesized in those cells, as with the Mendelian-like influence of *FUT2* genotype (i.e. secretor status) on certain HMOs^11^. Genetic variants that influence circulating levels of a component that is then transported into milk, for example glucose production in the liver, could also shape milk composition. Thus, identifying genetic factors that influence milk composition can help us better understand the biology of milk production and identify potential targets for intervention.

Although environmental and clinical influences on milk composition have been widely studied, genomic analyses of human milk remain limited. Most genome-wide association studies (GWAS) have focused on HMOs^12–14^, corroborating the known large effects at *FUT2*, with additional replicated associations at glycosyltransferases *FUT3* and *ST6GAL1*. A GWAS of milk fatty acids also identified associations at the fatty acid desaturase (*FADS*) genetic locus^15^, which had previously been described in candidate gene studies^16,17^. Overall, genetic studies of human milk lag far behind those of livestock, particularly dairy cattle, where large-scale GWAS have discovered numerous loci underlying traits such as milk yield and fat percentage^18^. Understanding the genetic basis of human milk composition will yield fundamental insights into the biology of milk production, enable genetic prediction of milk composition, and facilitate assessment of the health impacts of natural variation in milk composition.

Here, we describe a cross-sectional profile of the milk metabolome in 349 1-month human milk samples and comprehensively explore both genetic and non-genetic factors influencing the abundance of 458 milk metabolites. We identify dietary, genetic, transcriptomic, and clinical factors associated with variation in the milk metabolome. In metabolite GWAS, we find novel genetic associations with the milk metabolome. We then combine these factors with machine learning to identify the most informative maternal features shaping the abundance of milk metabolites. Overall, this work represents the first comprehensive analysis and comparison of genetic and non-genetic factors shaping the human milk metabolome, revealing new insights into the biology of human milk production.

## Results

### Metabolomic profiling in a human milk cohort

We explored human milk metabolomics data (n=349, **Figure 1A**, **Figure 1B, Supplementary Figure 1, Supplementary Table 1**) from the Mothers and Infants LinKed for Healthy Growth (MILK) study, an observational study of healthy full-term mothers and infants from Minnesota and Oklahoma, USA. Participants were primarily non-Hispanic white (77%) and had an average maternal age of 31.2 years, 27% had cesarean deliveries, and 15% had a gestational diabetes diagnosis (**Supplementary Table 2**). Utilizing a combination of three mass spectrometry strategies (Methods), semi-quantitative abundances of 458 milk metabolites were measured (**Figure 1C**). We identified seven modules of co-occurring metabolites (Methods; **Figure 1D, Supplementary Table 1, Supplementary Table 3**), and tested for enrichment of metabolite subclasses within each module (**Figure 1E, Supplementary Table 4**). Each module had at least one overrepresented metabolite subclass; for example, “fatty acid esters” were overrepresented in Module 1 (Fisher’s exact test, odds ratio = 26.5, q-value = 1.9×10^-12^), “amino acids, peptides, and analogues” were overrepresented in Module 2 (odds ratio = 11.3, q-value = 5.1×10^-9^). This indicates that co-occurring metabolites tended to belong to the same subclass.

**Figure 1.**
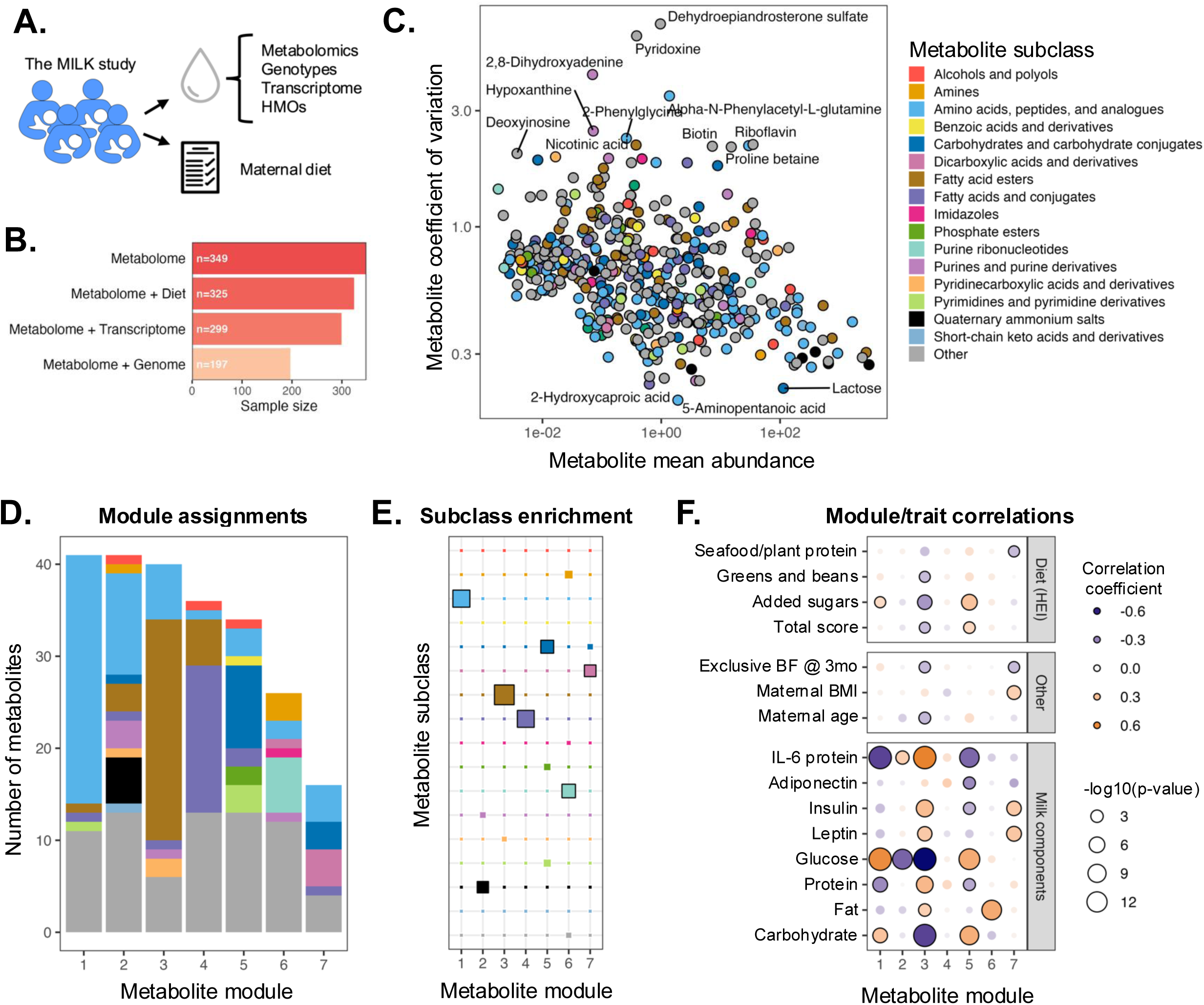
Overview of milk metabolome. **A)** Overview of data used in this manuscript. Milk samples from the MILK study were utilized to generate milk metabolomes, transcriptomes, genomes, and HMO profiles. Dietary patterns were assessed using the Dietary History Questionnaire (DHQ). **B)** Sample sizes for analyses described in this manuscript. **C)** Metabolite mean abundance and coefficient of variation. Each dot is a metabolite, colored by its metabolite subclass. **D)** Metabolites were assigned to seven modules of correlated metabolites using weighted gene co-expression network analysis (WGCNA). Metabolite subclasses are colored according to the legend in panel C. **E)** Enrichment of metabolite subclasses in each metabolite module as assessed by Fisher’s exact test. Tile size represents -log10(q-value). Module subclass pairs with significant enrichment (q-value < 0.05) are denoted with a black outline. Tiles are colored by metabolite subclass following the legend in panel C. **F)** Pairwise correlations between milk modules and traits. Circle color indicates Pearson correlation coefficient (purple = negative, orange = positive) and circle size indicates - log10(p-value). Significantly correlated module-trait pairs (q-value < 0.05) are denoted with a black circle outline. Maternal diet features are scores from the Healthy Eating Index (HEI).

To explore factors shaping the overall composition of the metabolome, we tested for correlations between each metabolite module, maternal factors including diet, and other milk components (**Figure 1F, Supplementary Table 5**). Here, non-metabolite milk components include macronutrient concentrations measured by transmission infrared spectroscopy, and hormones and cytokine concentrations measured by ELISA (Methods). We observed stronger correlations between the milk metabolome and other milk traits than between the metabolome and maternal traits (**Figure 1F**). The strongest correlations appeared to capture a signature of inflammation described in our previous study, where we found that milk IL-6 protein concentration was the factor that explained the most variation in the milk transcriptome^19^. Here, milk samples with higher concentrations of IL-6 protein and lower concentrations of glucose tended to have higher abundances of the fatty acid esters in Module 3 (**Supplementary Figure 2**). These fatty acid esters are acylcarnitines, metabolites that transport fatty acids into the mitochondria to generate ATP via β-oxidation, and included short, medium, and long chain acylcarnitines (**Supplementary Figure 3**). Among maternal non-milk traits, we observed the strongest relationships between maternal diet and the milk metabolome (**Supplementary Table 5**). Overall, this analysis identified multiple maternal factors that shape broad patterns of the milk metabolome.

### Dietary influences on the milk metabolome

As many milk metabolites are derived directly or indirectly from maternal nutrient intake, we examined relationships between maternal diet and milk metabolites to gain insight into modifiable exposures that shape the nutritional and bioactive properties of human milk. Maternal diet patterns were measured with the Diet History Questionnaire (DHQ, Methods), a food frequency questionnaire that quantifies intake of specific nutrients and food categories, in 344 participants with milk metabolomics data.

We first tested for pairwise correlations between diet features and milk metabolite abundances. Comparing 199 DHQ-derived dietary features to 458 metabolites, we identified 1,038 pairs with significant correlations (Kendall’s *τ* q-value < 0.05; **Supplementary Table 6**). Of these significant diet-metabolite pairs, 920 (88%) were positive correlations (**Supplementary Figure 4**), likely reflecting higher dietary intake resulting in higher levels of a nutrient or its byproducts in breast milk. The “fatty acid esters” metabolite subclass was an exception, with 36/38 pairwise correlations being negative (**Supplementary Figure 4, Supplementary Table 6**). This observation suggests diet may influence milk acylcarnitines more through a change in metabolic state rather than as direct food-derived components.

To identify the maternal diet features that best explain the milk metabolome, we performed lasso feature selection (Methods) from DHQ-derived dietary features (**Figure 2A; Supplementary Table 7**). To reduce collinearity, we pruned the 199 DHQ variables by iteratively removing one variable from a pair with Pearson correlation coefficient > 0.99, resulting in 174 DHQ variables available for feature selection. Lasso regression identified features with non-zero regression coefficients for 371 of 458 metabolites (**Figure 2A**, **Supplementary Table 7**). The selected dietary features explained a median of 1.2% of variation in each metabolite, with a median of 6 dietary features selected per metabolite.

**Figure 2.**
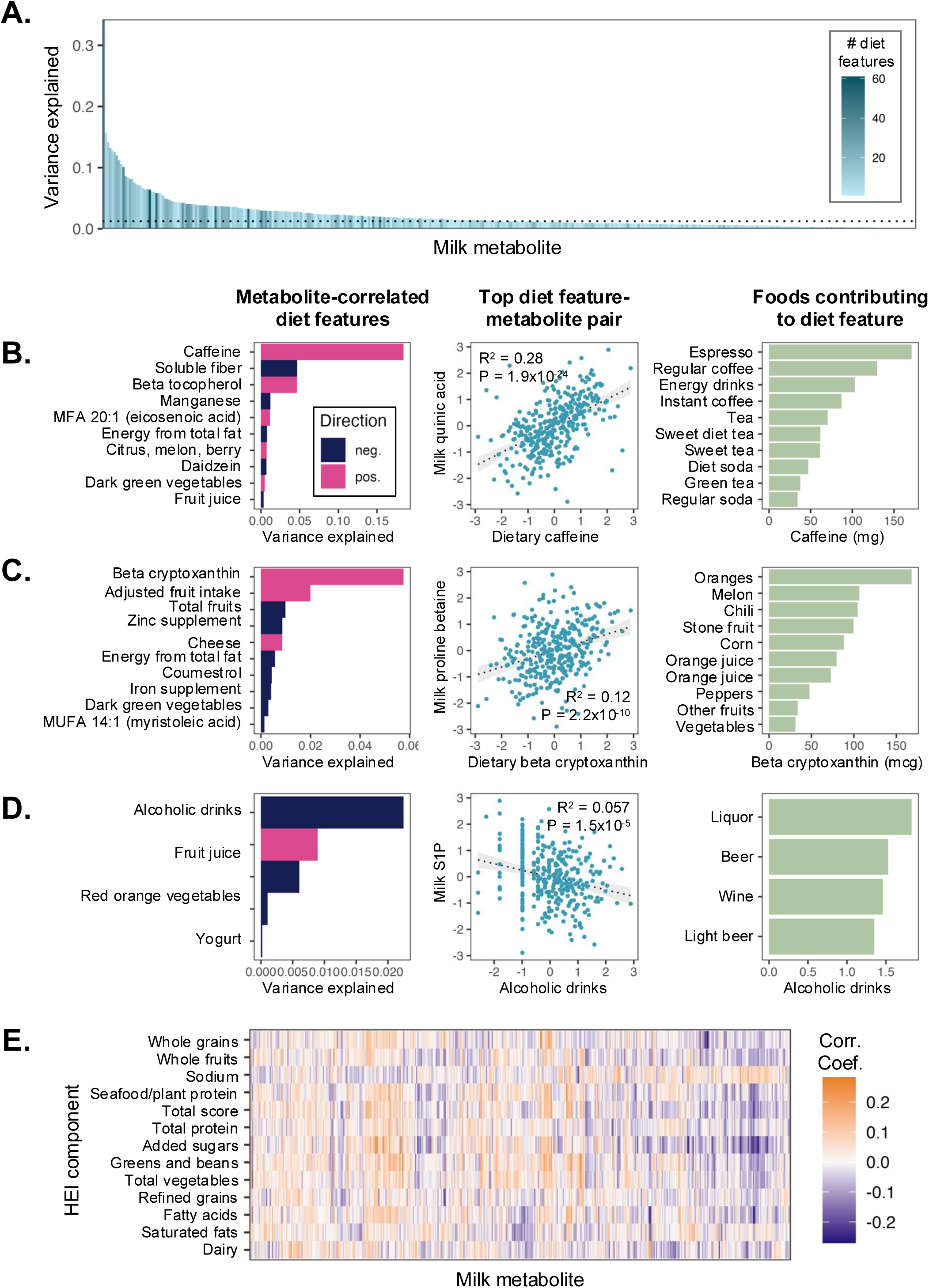
Dietary influences on the human milk metabolome. **A)** Variation explained in each milk metabolite (out-of-fold R^2^) by the dietary features selected by lasso regression model. Each vertical bar is a metabolite. Bar color represents the number of dietary features selected by the model. The dotted horizontal line represents the median R^2^. **B-D)** Characteristics of dietary features contributing to (B) quinic acid, (C) proline betaine, and (D) S1P (sphingosine-1-phosphate). Left panel: top dietary features selected for each metabolite in milk, bar color indicating direction of correlation with metabolite abundance. Dark purple bars indicate negative correlation with the milk metabolite, magenta bars indicate positive correlation with the milk metabolite. Middle panel: scatter plot of normalized metabolite abundance (y-axis) and normalized dietary features (x-axis), with each point representing a milk sample. R² and P values in the middle panels are from the displayed pairwise associations and are distinct from the out-of-fold lasso model R² shown in the left panels. Right panel: foods contributing most to the DHQ dietary feature in middle panel. **E)** Pearson correlations between milk metabolite abundances and Healthy Eating Index (HEI) component scores. Metabolites and HEI component scores are arranged by hierarchical clustering.

Examining the metabolites best explained by dietary features, quinic acid stands out with 34% variance explained (**Figure 2B**), due to its strong relationship with caffeine consumption. This was also the strongest pairwise correlation between any milk metabolite and dietary feature (Kendall’s *τ* = 0.41, q-value = 6.9×10^-21^). Caffeine intake is one of the foods most accurately measured by questionnaire^20^, as people often routinely drink the same number of cups of coffee or tea per day and abstainers will accurately report their lack of coffee or tea. Thus, the exceptionally strong correlation between dietary caffeine and coffee-derived metabolites compared to other foods may be in part due to the relative accuracy of the questionnaire data for caffeine.

Proline-betaine was also well-explained by dietary features, with 12.5% variance explained (**Figure 2C**). The most predictive selected dietary feature for proline-betaine (a.k.a. stachydrine) was beta cryptoxanthin. Citrus fruits are among the foods with the highest levels of beta cryptoxanthin^21^, and proline-betaine is abundant in citrus fruits and has been validated as a biomarker of citrus consumption^22^. Another diet-correlated metabolite was sphingosine-1-phosphate (S1P; R^2^ = 0.038). The top-weighted dietary feature selected for S1P was number of alcoholic drinks, with higher alcohol intake correlated with lower levels of S1P in milk (**Figure 2D**). A recent report showed S1P decreased in mice after alcohol injection^23^. S1P is a signaling lipid that may have roles in infant gut maturation and immune maturation^24^.

We also compared the milk metabolome to overall diet quality. DHQ data was summarized by the Healthy Eating Index^25,26^ (HEI), comprised of 13 component scores and a total score that quantify adherence to dietary guidelines, with higher scores indicating better diet quality. Examining pairwise correlations between milk metabolites and HEI scores (**Figure 2E, Supplementary Table 8**), HEI total score was positively correlated with milk riboflavin and N-acetylornithine (**Supplementary Figure 5**). We next used lasso regression to determine if milk metabolites could predict overall diet quality as assessed by HEI scores (Methods, **Supplementary Table 9**). Lasso selected metabolite features for 13 of 14 HEI components, including total HEI score but excluding the “refined grains” index. The “greens and beans” index, quantifying intake of dark green vegetables and legumes, was the HEI component best explained by the milk metabolite data (out-of-fold R^2^ = 0.09 (**Supplementary Figure 6**).

Overall, our exploration of dietary patterns and the milk metabolome identified correlated milk metabolites for the majority of diet features as well as overall dietary patterns. These results underline the influence of maternal diet on milk composition and suggest milk as a potential tool to profile maternal diet patterns.

### Associations between the milk metabolome and milk cell transcriptome

One strength of our cohort is deep multi-omic profiling of milk, including 299 1-month samples with both milk metabolome and bulk milk cell transcriptome data. To identify the most closely linked gene-metabolite pairs, we used lasso regression (Methods). Lasso identified gene features with non-zero weights for 398 metabolites (of 458), with a median of 20 genes selected per metabolite (**Figure 3A, Supplementary Table 10**). The selected features explained a median of 2.8% of variation in each metabolite abundance (**Figure 3A**). The best-explained metabolite was hydroxybutyrylcarnitine, a short-chain acylcarnitine (R^2^ = 0.30).

**Figure 3.**
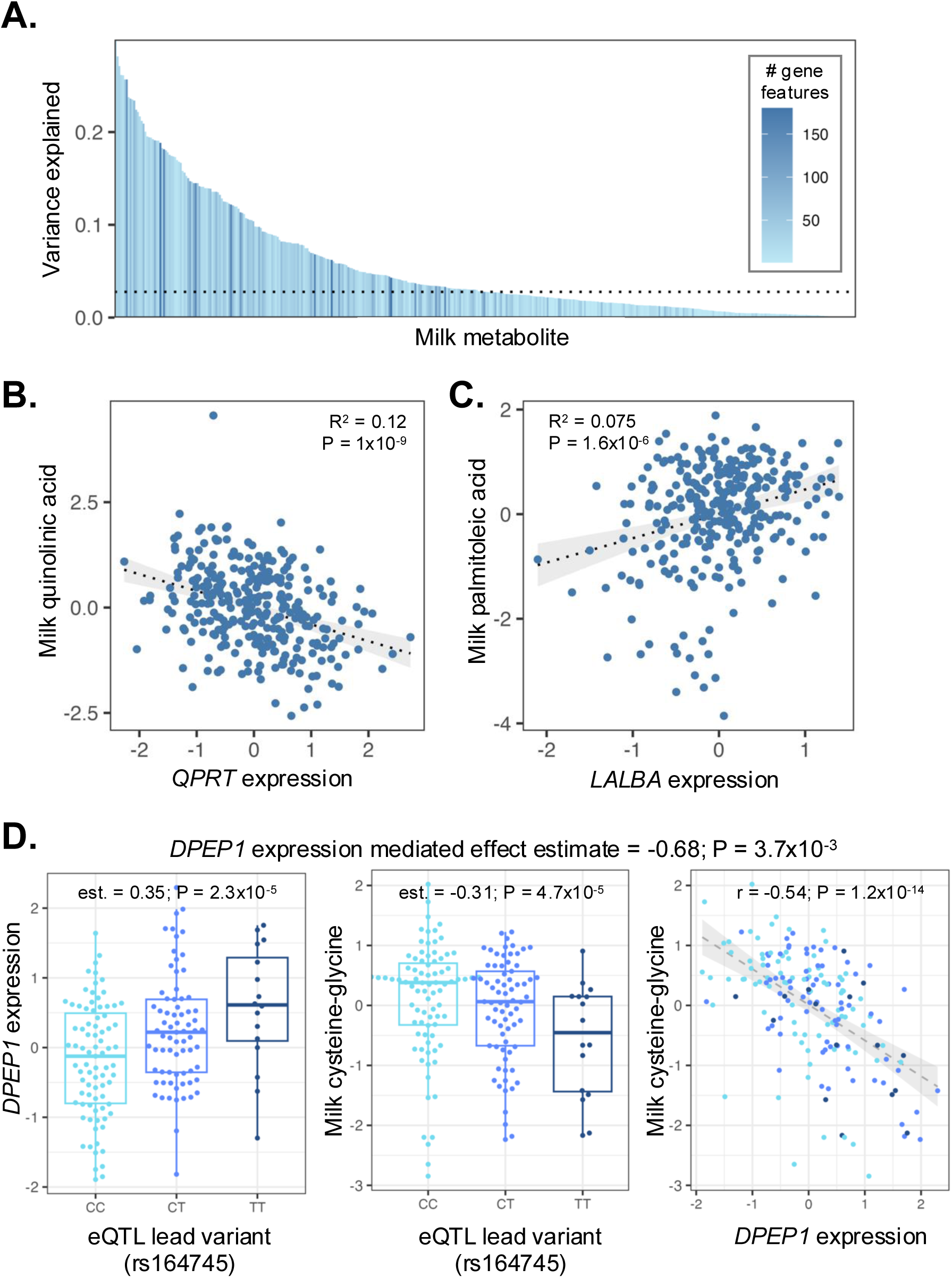
Relationships between human milk cell gene expression and the milk metabolome. **A)** Variation explained in each milk metabolite (out-of-fold R^2^) by the milk-expressed genes selected by lasso regression model. Each vertical bar is a metabolite, and bar color represents the number of genes selected by the model. The dotted horizontal line represents the median R^2^. **B)** Scatter plot of normalized milk quinolinic acid abundance and *QPRT* expression. R^2^ and p-value from the pairwise correlation of gene expression with metabolite. **C)** Scatter plot of normalized milk palmitoleic acid abundance and *LALBA* expression. R^2^ and p-value from pairwise correlation of gene expression with metabolite. **D)** Relationships between the lead milk eQTL variant for *DPEP1* (rs164745) and milk cysteine-glycine abundance (left panel), *DPEP1* expression (middle panel); and scatter plot of *DPEP1* expression and cysteine-glycine (right panel).

Overall, lasso feature selection identified relevant genes for milk metabolites. For example, the top gene feature for quinolinic acid was *QPRT* (quinolinate phosphoribosyltransferase) (**Figure 3B**; partial R^2^ = 0.097, p-value = 1.0×10^-7^), with a negative correlation reflecting that enzyme’s role in converting quinolinic acid into nicotinic acid mononucleotide. This analysis also highlighted the relationship between expression of tryptophan-to-kynurenine enzyme *IDO1* in shaping milk levels of kynurenine (partial R^2^ = 0.080, p-value = 3.1×10^-6^), a pathway we previously discovered is upregulated in the presence of cytomegalovirus in human milk^27^. Another interesting example was *LALBA*, the highly expressed milk gene encoding the regulatory subunit of lactose synthase, as the top gene feature for fatty acids alpha-linolenic acid and palmitoleic acid (**Figure 3C**; partial R^2^ = 0.082, p-value = 8.8×10^-7^). Alpha-linolenic acid is an essential omega-3 fatty acid that cannot be synthesized *de novo* in humans and therefore must be obtained from the diet^28^, whereas palmitoleic acid, an omega-7 monounsaturated fatty acid, can be synthesized endogenously from palmitic acid through stearoyl-CoA desaturase^29^. Thus, the relationship between *LALBA* and these fatty acids, both common in human milk, likely reflects overall milk synthetic state in the mammary gland.

The top gene feature for the dipeptide cysteine-glycine was *DPEP1* (dipeptidase 1) (**Figure 3D)**. This enzyme is expressed across luminal membrane tissues including lactating mammary gland^30^ and hydrolyzes dipeptides including cysteine-glycine. As *DPEP1* has a milk expression quantitative trait locus (eQTL)^19^, we tested for a causal relationship between *DPEP1* gene expression and cysteine-glycine in our milk samples using two-stage least squares regression analysis (Methods). We found evidence for a causal relationship, with each standard deviation increase in *DPEP1* expression associated with a -0.68 standard deviation decrease in cysteine-glycine in milk (p-value = 3.7×10^-3^). Colocalization also supported this interpretation, with a posterior probability of shared causal variant of 0.84 (**Supplementary Figure 7**). The lead variant for the milk eQTL (rs164745, chr16:89643256:C:T) is also associated with plasma cysteine-glycine levels^31^, *DPEP1* expression in kidney, kidney function, and risk for kidney disease^32^; suggesting that its link with milk cysteine-glycine levels could be due to *DPEP1* expression in other tissues if the dipeptide is transported from circulation into lactocytes^33^. Cysteine-glycine provides an amino acid source to the infant, and is also a precursor and product of glutathione, which functions in the antioxidant system of lactating mammary gland^34^.

### GWAS of milk metabolites identifies novel metabolite QTL unique to human milk

Motivated by the biologically plausible gene-metabolite connections identified in our comparison of milk metabolomes and transcriptomes, we next performed a GWAS for each milk metabolite in n=197 unrelated samples with both genotype and milk metabolomics data. While this is a relatively small sample size for a GWAS, because metabolite levels are biologically closer to the effects of genetic variation than more complex organismal phenotypes, genuine genetic associations can often be identified in cohorts of only a few hundred individuals^35^.

There were nine study-wide significant locus-metabolite pairs, at five distinct loci (p-value < 1.09×10^-10^, Bonferroni correction of 5×10^-8^ / 458 metabolites) (**Table 1**, **Figure 4A, Supplementary Table 11**). Two of these loci (*PDE6A* and *GNE*) represent new metabolite associations, discussed below. The association with imidazoleacetic acid riboside near the *NAPRT* gene, a ribosyl transferase gene on chromosome 8, has been previously described in cerebrospinal fluid and plasma^36,37^. This locus is also an eQTL for *NAPRT* in human milk, and mediation analysis in our data supported a causal role for *NAPRT* gene expression in shaping imidazoleacetic acid riboside levels in milk (**Figure 4B**, Methods). Additional study-wide significant metabolite associations that have previously been described in other tissues include N-acetylornithine levels (and N-alpha-acetylarginine at genome-wide significance, p = 1.45×10^-8^) near *NAT8*, an N-acetyltransferase gene on chromosome 8; and S-adenosyl-L-methionine near *SLC25A26*, a mitochondrial S-adenosyl-L-methionine antiporter on chromosome 3. The association of free fucose monosaccharide with variants in *FUT2*, the fucosyltransferase gene that confers secretor status on chromosome 19, has previously been reported in milk as an association between secretor status and free fucose^38^.

**Figure 4.**
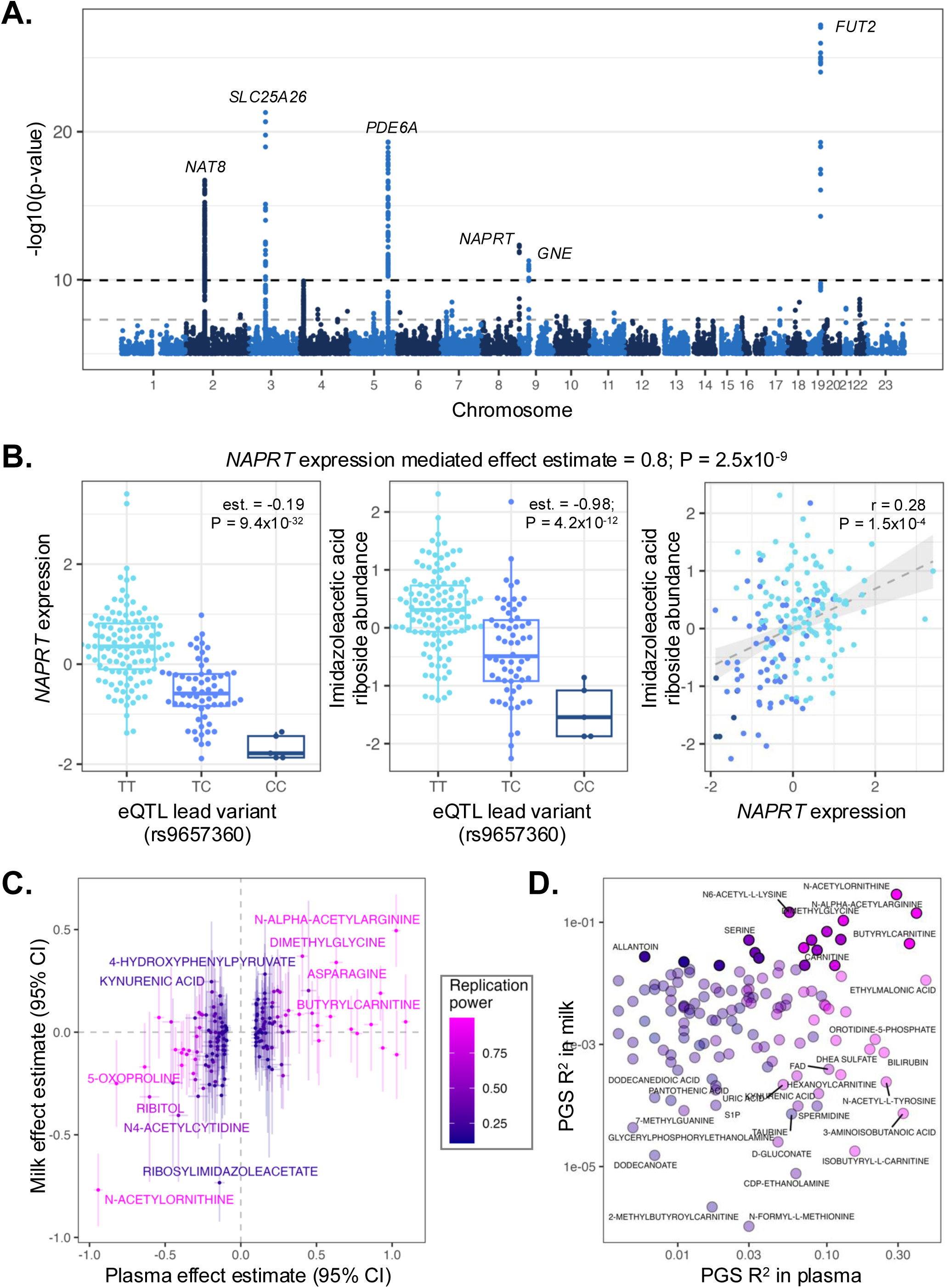
GWAS of human milk metabolome. **A)** Manhattan plot of GWAS of all 458 milk metabolites. Only variants with p-value < 5×10^-5^ are plotted. Gene names are displayed for loci with study-wide significant associations (p-value < 1.09×10^-10^). **B)** Relationships between the lead milk eQTL variant for *NAPRT* (rs9657360) and *NAPRT* expression (left panel), milk imidazoleacetic acid riboside abundance (middle panel); and scatter plot of *NAPRT* expression and imidazoleacetic acid riboside (right panel). **C)** Scatter plot of plasma metabolite QTL replication in milk. Each dot is a study-wide significant metabolite genetic association from Chen et al., with the x-axis plotting the plasma effect estimate and the y-axis plotting the milk effect estimate, with error bars for 95% confidence intervals. Points are colored by the estimated power to replicate the association in plasma in our milk dataset. **D)** Scatter plot of variance explained in plasma metabolites (x-axis) and milk metabolites (y-axis) by a PGS derived from plasma genetic association data. Points are colored by the estimated power to replicate the association in plasma in our milk dataset.

**Table 1.** Study-wide significant loci.

| **Metabolite** | **rsID** | **Variant ID** | **P-value** | **Gene** | **Gene function** |
|---|---|---|---|---|---|
| N-acetylornithine | rs35541168 | 2:73621272:CT:C | 1.90x10<sup>-17</sup> | *NAT8* | N-acetyltransferase |
| S-adenosyl-L-methionine | rs1084535 | 3:66309606:G:A | 5.03x10<sup>-22</sup> | *SLC25A26* | Mitochondrial S-adenosyl-L-methionine antiporter |
| Hypoxanthine | rs11960069 | 5:149929538:T:C | 2.07x10<sup>-12</sup> | *PDE6A* | Phosphodiesterase that catalyzes hydrolysis of 3',5'-cyclic GMP |
| AICA riboside | rs1991805 | 5:149931649:G:C | 6.07x10<sup>-14</sup> | *PDE6A* | Phosphodiesterase that catalyzes hydrolysis of 3',5'-cyclic GMP |
| Xanthine | rs1991805 | 5:149931649:G:C | 2.12x10<sup>-17</sup> | *PDE6A* | Phosphodiesterase that catalyzes hydrolysis of 3',5'-cyclic GMP |
| N-formyl-GAR | rs17725010 | 5:149933781:C:T | 4.98x10<sup>-20</sup> | *PDE6A* | Phosphodiesterase that catalyzes hydrolysis of 3',5'-cyclic GMP |
| Imidazoleacetic acid riboside | rs10093709 | 8:143602045:A:G | 4.62x10<sup>-13</sup> | *NAPRT* | Ribosyl transferase |
| N-acetyl-neuraminic acid (Neu5Ac) | rs10972852 | 9:36319084:G:T | 5.22x10<sup>-12</sup> | *GNE* | Regulates and initiates biosynthesis of N-acetylneuraminic acid |
| Fucose | rs2251034 | 19:48704535:A:G | 6.16x10<sup>-28</sup> | *FUT2* | Fucosyltransferase |

Given that several of the milk metabolite quantitative traitl loci (mQTL) we identified were previously observed in larger plasma mQTL studies, we explored the extent to which plasma mQTL replicate in milk. We identified 152 mQTL that were study-wide significant (p-value < 4.6×10^-11^) in a plasma metabolite GWAS of 8,299 Canadians^37^ and had the same metabolite and lead genetic variant available in our milk dataset (**Supplementary Table 12**). Overall, effect estimates between plasma and milk were correlated (R^2^ = 0.22, p-value = 9.0×10^-10^; **Figure 4C**). Though lack of replication cannot prove the absence of a genetic effect on milk metabolites, we observed some interesting non-replicated examples (milk mQTL p-value > 0.2) with >90% power to replicate the genetic effect seen in plasma (**Supplementary Figure 8**). Three of seven non-replicating loci (plasma mQTL at *ACADM*, *ACADS*, and *SLCO1B1*) involve fatty acid metabolism, highlighting differences in lipid metabolism between lactating mammary gland and plasma. We also noted that while the *NAT8* locus association with acetylated amino acids replicated for acetylornithine and acetylarginine, it did not replicate for glutamate-family amino acids, including acetylglutamate, acetylglutamine, and acetylasparagine (**Supplementary Figure 8**, **Supplementary Table 12**). Glutamate is ∼40x more abundant in human milk relative to plasma^39,40^, likely due to substantial de novo synthesis within the mammary gland^40^; thus a weak relationship between genetic effects on plasma (acetyl)glutamate levels and those in milk is plausible.

We also explored whether polygenic scores (PGS) derived from plasma metabolite GWAS^41^ predicted metabolite abundances in our milk dataset (**Figure 4D**, **Supplementary Table 13**). Of 135 milk metabolites with a plasma PGS, 19 had a nominally significant correlation between PGS and milk metabolite abundance (p-value < 0.05) (**Figure 4D**). For all metabolites with a significant correlation with the plasma PGS, higher PGS corresponded to increased milk metabolite abundance (**Supplementary Figure 9**), indicating broad concordance despite our overall lack of power due to limited sample size. The best-predicted metabolite was N-acetylornithine (milk R^2^ = 0.29, p-value = 6.0×10-16), primarily due to the large influence of the *NAT8* mQTL locus variants included in the plasma PGS (**Supplementary Figure 10**).

We observed a locus on chromosome 5 that had study-wide significant associations with four metabolites (AICA riboside, N-formyl-GAR, hypoxanthine, and xanthine; **Table 1**) and a genome-wide significant association with acetylphosphate (p-value < 5×10^-8^; **Supplementary Figure 11**). All study-wide significant metabolites at this locus are part of the purine metabolism pathway (**Figure 5A**). The fine-mapped credible sets for each metabolite fall within *PDE6A* on chromosome 5 (**Supplementary Figure 12**), which encodes a phosphodiesterase that catalyzes the hydrolysis of 3’,5’-cyclic GMP (**Figure 5A**). The directions of SNP effects are consistent with this pathway, with metabolites upstream of GMP having positive effect estimates and downstream metabolites having negative SNP effect estimates. *PDE6A* has mostly been studied for its role in phototransduction in rods and cones^42^, but is also expressed in other tissues including pituitary, adrenal and salivary glands^43^ (**Supplementary Figure 13**). Genetic effects of the same SNP on *PDE6A* gene expression were identified in esophagus mucosa and pituitary tissues in GTEx (**Supplementary Figure 14**)^43^. While *PDE6A* did not pass the minimum expression threshold to be included in our eQTL study of human milk^19^, the direction of effect on *PDE6A* expression of the mQTL lead SNP is consistent with that in esophagus mucosa (**Supplementary Figure 14**). This locus also contains a milk eQTL for *SLC26A2*, a nearby sulfate transporter gene (**Supplementary Figure 14**). As expected, abundances of metabolites within this pathway are intercorrelated (**Supplementary Figure 15**). We explored the colocalization patterns of the metabolite QTL and expression QTL for *PDE6A* and *SLC26A2* and found that all the milk mQTL colocalize with each other and with the *PDE6A* eQTL in esophagus and pituitary gland (posterior probability of shared causal variant > 0.9; **Supplementary Figure 16**, **Supplementary Table 14**). The lead variant at this milk mQTL (rs17725010) has no significant association for xanthine or hypoxanthine (p>0.05) in plasma^44^. Given the combined genetic and metabolic pathway evidence, we hypothesize that this locus shapes the abundance of purine pathway metabolites via *PDE6A* expression in a milk-specific manner. Hypoxanthine and xanthine in milk, in combination with the highly-expressed xanthine oxidase enzyme, have an antimicrobial effect^45^, suggesting genetic variation at this locus could impact the innate immune properties of human milk.

**Figure 5.**
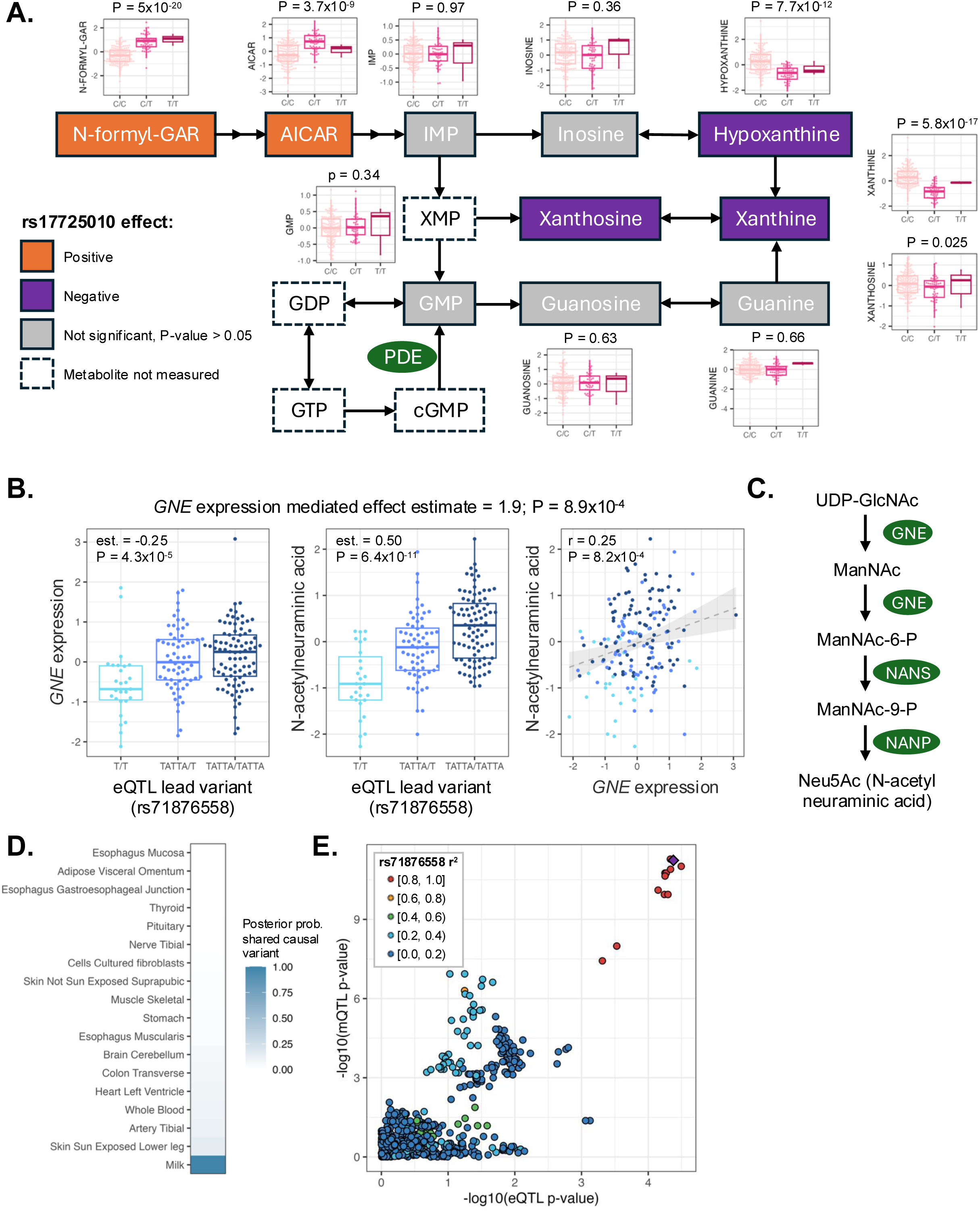
Novel milk-specific metabolite QTL at the *PDE6A* and *GNE* loci. **A)** Diagram of metabolites in the purine metabolism pathway. Boxplots alongside each metabolite display genetic associations with the lead variant at this locus (rs17725010). PDE (phosphodiesterase) enzymes (green oval), including PDE6A, catalyze the hydrolysis of 3’,5’-cyclic GMP into GMP. Pathway metabolite boxes are colored orange for positive genetic effects, purple for negative genetic effects, gray for non-significant effect (p-value > 0.05), or white with a dashed outline if the metabolite was not measured in our dataset. **B)** Relationships between the lead milk eQTL variant for *GNE* (rs71876558) and *GNE* expression (left panel), N-acetyl neuraminic acid (Neu5Ac) (middle panel); and scatter plot of *GNE* expression and Neu5Ac in milk (right panel). **C)** Diagram of Neu5Ac biosynthesis pathway. GNE initiates and regulates the Neu5Ac biosynthetic pathway. **D)** Colocalization of milk Neu5Ac QTL with *GNE* eQTL in milk (this cohort) or other tissues (GTEx). **E)** Scatter plot of GNE eQTL p-values (x-axis) and Neu5Ac QTL (y-axis) in milk.

We identified a second potentially milk-specific mQTL for N-acetyl-neuraminic acid (Neu5Ac) on chromosome 9 upstream of *GNE* (**Figure 5B**, p-value = 5.2×10^-12^), which encodes the rate-limiting enzyme in Neu5Ac biosynthesis (**Figure 5C**). Neu5Ac is the predominant form of sialic acid in humans^46^. Mediation analysis supported a causal relationship of *GNE* expression mediating the genetic association with milk Neu5Ac levels (**Figure 5B**, Methods). The Neu5Ac mQTL colocalizes with a *GNE* eQTL in milk (**Figure 5D, 5E**; posterior probability of shared causal variant = 0.99) but not with *GNE* eQTL for any other GTEx tissue (**Figure 5D**). *GNE* is widely expressed across tissues^43^, with milk gene expression falling in the mid-range of other tissues (**Supplementary Figure 17**). The lead variant for the milk mQTL at the *GNE* locus (rs71876558) was not associated with Neu5Ac in larger metabolite GWAS of cerebrospinal fluid^47^ or plasma^37^ (p-values > 0.05). Neu5Ac abundance in milk was uncorrelated with a plasma PGS for Neu5Ac^41^ (**Supplementary Figure 18**; R^2^ = 7×10^-4^, p-value = 0.71) that included variants at 2 loci near genes relevant to sialic acid biology (*NPL*, *ST5GALNAC1*) but not *GNE*. Thus, the available evidence suggests this association is unique to sialic acid biosynthesis in the lactating mammary gland.

Human milk is among the tissues with highest concentrations of sialic acid, along with the brain and central nervous system^46^. In human milk, a majority of the sialic acid is bound in human milk oligosaccharides (HMOs) or proteins, with only 3% as free sialic acid^48^. The mQTL identified here is an association with free sialic acid, but we have also measured HMO concentrations in our samples. Thus, we tested if the genetic variant associated with free sialic acid also associates with the concentrations of sialylated HMOs. We did not observe an association between the sialic acid mQTL lead variant and the 19 HMOs measured in our study, but there was a nominal association with the sialylated HMO LSTb in a larger HMO GWAS^13^ (**Supplementary Figure 19**). These results suggest that the effect of the mQTL on free sialic acid seen here does not translate into a large effect on HMO composition.

### Integration of maternal genetic and non-genetic factors shaping the milk metabolome

Finally, we combined maternal genetic, dietary, and clinical data to explore the relative contributions of these factors in shaping the milk metabolome. We used lasso feature selection to identify the most predictive features for each metabolite (**Figure 6A**). Lead variants of mQTL that passed genome-wide significance (p-value < 5×10^-8^, **Supplementary Table 11**) and all diet and clinical features were included as potential features. We excluded milk gene expression and other milk composition measurements from this analysis, seeking to understand influences most likely upstream of the milk metabolome. We note that the variation explained by genetics here represents just a small number of genetic variants (i.e., mQTL lead variants), rather than an estimate of genome-wide SNP heritability, which would require a much larger dataset. The lasso model selected features for 246 out of 458 metabolites (**Supplementary Table 15**). These models explained a median of 13.2% of metabolite variation, with diet features explaining the most variation in 168 (68%) of metabolite models (**Figure 6A**).

**Figure 6.**
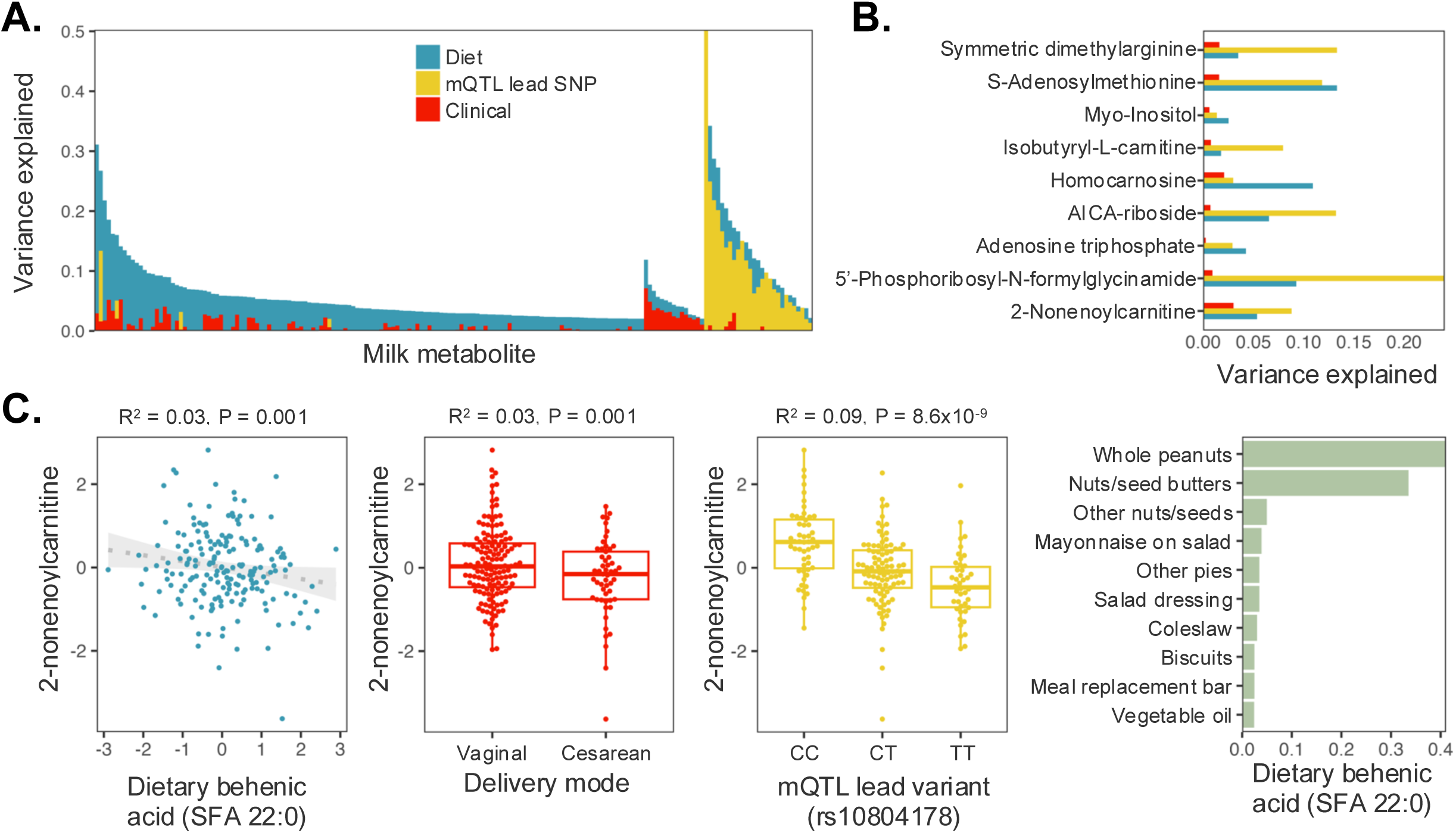
Relative contributions of dietary and genetic influences to human milk metabolites. **A)** Variation explained in each milk metabolite (out-of-fold R^2^) by the maternal genetic, dietary, and clinical features selected by lasso regression model. Each vertical bar is a metabolite, and bar height represents the proportion of variance explained by the model for each metabolite. Bar colors represent the proportion of variation explained by each feature type. B) For metabolites with features selected from all three categories (diet, genetics, and clinical), the proportion of variation explained by each feature type. **C-D)** Characteristics of features contributing to 2-nonenoylcarnitine (C) and N-acetylornithine (D). Left panel: scatter plot of top diet feature vs. metabolite abundance. Middle left panel: Boxplot of top clinical feature vs. metabolite abundance. Middle right panel: Boxplot of top genetic variant vs. metabolite abundance. Right panel: foods contributing most to the top dietary feature.

There were 12 metabolites for which features from all three data types (dietary, clinical, genetic) were selected (**Figure 6B**). For example, for 2-nonenoylcarnitine, lasso selected six features, with the top feature from each category plotted in **Figure 6C**. 2-nonenoylcarnitine is an acylcarnitine comprised of a medium-chain unsaturated fatty acid (C9:1) esterified to carnitine for transport into mitochondria for β-oxidation. The top overall feature was a genome-wide significant mQTL (lead SNP: rs10804178, mQTL p-value = 2.3×10^-8^) that explained 18% of variation in 2-nonenoylcarnitine abundance in milk. This mQTL resides in *UNC80* and is ∼240kbp downstream of *ACADL*, which encodes an acyl-CoA dehydrogenase that catalyzes the first step of mitochondrial fatty acid β-oxidation. This locus is associated with nonanoylcarnitine (saturated C9 acylcarnitine) in human plasma (p-value = 2.05×10^-234^)^44,49,50^. The top dietary feature was SFA 22:0 (behenic acid), a very-long-chain saturated fatty acid found in plant oils including peanut oil, which explained 4.6% of variation in 2-nonenoylcarnitine. Higher behenic acid consumption correlated with lower 2-nonenoylcarnitine in milk. Among clinical features, cesarean delivery explained an additional 6.1% of variation, with women who delivered via cesarean having lower levels of 2-nonenoylcarnitine in their milk.

Another metabolite with selected features from all 3 categories was N-acetylornithine (**Figure 6D**), with its milk mQTL as its top feature (rs35541168, 29% of variation in milk metabolite levels, mQTL p-value = 1.9×10^-17^). Additional selected features for milk N-acetylornithine included a positive correlation with dietary coumestrol (3.8% of variation) and a negative correlation with gestational diabetes (1.8% of variation). Coumestrol is a phytoestrogen found primarily in legumes^51^. In a mouse study comparing milk metabolites after feeding obesogenic vs. standard diets, N-acetylornithine was one of three highly discriminant milk metabolites between diets (along with proline-betaine and TMAO)^52^. N-acetylornithine and proline-betaine have also been identified as biomarkers of a healthy diet in plasma^53^. Similarly, we observed a positive correlation between diet features associated with a healthier, high-fiber diet and these milk metabolites (e.g. citrus fruit and proline-betaine, **Figure 2C**; nuts/seeds and N-acetylornithine, **Figure 6D**). TMAO was also quantified in our dataset, and its top selected feature was a positive correlation with dietary docosapentaenoic acid (R^2^ = 0.099, p-value = 1.3×10^-5^), a polyunsaturated fatty acid found in oily fish like salmon^54^ (**Supplementary Figure 20**). TMAO (trimethylamine-N-oxide) is primarily produced from dietary precursors by microbes in the gut and is positively correlated with heart and kidney disease^55^. However, in our dataset as in the mouse study^52^, TMAO in milk was positively associated with healthier diet features.

## Discussion

This work provides the first integrated analysis comparing clinical, dietary, transcriptomic, and genomic influences on the human milk metabolome. Our findings demonstrate that maternal diet broadly shapes milk metabolite composition, while a subset of metabolites exhibit strong genetic regulation. In the first milk metabolome GWAS, we identified two novel loci not previously observed in other human tissues. More broadly, these results establish the human milk metabolome as a rich molecular phenotype that captures both maternal exposures and mammary gland biology. Future longitudinal and mechanistic studies will help clarify how variation in milk metabolites influences infant development and whether specific milk metabolites can serve as biomarkers or targets for nutritional intervention during lactation.

We observed a positive correlation between acylcarnitines and milk IL-6 concentrations. In our previous analysis of this cohort, IL-6 explained more variation in the milk transcriptome than any other measured milk component^19^. Similarly, an increase in plasma and milk acylcarnitines was previously observed in dairy cattle following a lipopolysaccharide challenge to trigger an inflammatory response^56^. Acylcarnitines transport fatty acids into mitochondria for β-oxidation. Although the lactating mammary gland is highly lipogenic^10^, increased acylcarnitine abundance may indicate a shift toward greater mitochondrial fatty acid oxidation under inflammatory or metabolically stressed conditions^57^. Alternatively, elevated acylcarnitines may reflect changes in milk cellular composition accompanying inflammation. Distinguishing these possibilities will require future studies combining single-cell transcriptomics with metabolic measurements in human milk. Notably, acylcarnitines comprised the only metabolite subclass with predominantly negative associations with dietary variables, suggesting they may reflect maternal metabolic state rather than direct dietary transfer.

Though previous smaller human studies (n<40) found weak evidence for links between maternal diet and the milk metabolome^2,58,59^, our analyses revealed extensive links between maternal dietary patterns and milk metabolites. Many of the strongest associations reflected biologically plausible relationships between dietary intake and known food-derived metabolites, including associations between caffeine intake and coffee-derived metabolites such as quinic acid and 1,3-dimethyluric acid, and between citrus-associated dietary features and proline-betaine. We also observed a negative correlation between alcohol intake and sphingosine-1-phosphate, consistent with alcohol’s impact on sphingosine signaling in other tissues^23,60^. S1P is a bioactive sphingolipid involved in epithelial barrier function and intestinal development^61^, making it an intriguing candidate mediator linking maternal diet with infant gut biology. These findings substantially expand the number of diet-associated milk metabolites in human milk. Importantly, diet-derived features were selected by lasso regression for the majority of milk metabolites, suggesting that maternal nutrition exerts broad influence on metabolic pathways active during lactation. Milk metabolites showing the strongest relationships with dietary patterns (i.e. N-acetylornithine and proline-betaine) were also highlighted in prior studies of human plasma^62^ and mouse milk^52^, suggesting that some milk metabolites serve as reproducible biomarkers of maternal dietary quality across species. Overall, our results highlight diet as a key modifiable factor shaping human milk, and position milk metabolomics as a non-invasive tool to quantify maternal nutritional status in conjunction with subjective dietary surveys. Furthermore, future studies should investigate whether diet-modifiable metabolites such as N-acetylornithine and proline-betaine influence infant health outcomes, potentially informing dietary recommendations during lactation.

The strong influence of dietary caffeine on the milk metabolome raises the question of what impact this variation has on infant health and development. In particular, coffee influences the adult gut microbiome, specifically by driving an increase in the bacterial species *Lawsonibacter asaccharolyticus* which covaries with plasma quinic acid and other coffee-derived metabolites^63^. Whether the coffee-linked milk metabolites identified here also influence infant *L. asaccharolyticus,* and more generally the effects of other diet-linked milk metabolites on the infant gut microbiome, will be the focus of future work.

Our genome-wide association analyses replicated known metabolite quantitative trait loci and identified new putative milk-specific effects. The strongest examples of milk-specific regulation involved loci near *PDE6A* and *GNE*. At the *PDE6A* locus, multiple metabolites within the purine metabolism pathway shared colocalizing genetic signals, and the direction of effects across metabolites was consistent with pathway structure. Similarly, the association between variants near *GNE* and free N-acetyl-neuraminic acid abundance in milk colocalized with a milk eQTL but not with eQTL in other GTEx tissues, supporting tissue-specific regulation of sialic acid metabolism in the lactating mammary gland. Because sialic acid is highly abundant in human milk and important for infant neurodevelopment and immune function, this result points toward potentially unique genetic regulation of mammary gland glycan biology.

Our study’s strengths include deep molecular profiling of a well-characterized lactation cohort with paired metabolomics, transcriptomics, genotypes, and dietary data. We acknowledge several limitations: first, all milk was sampled at approximately one month postpartum, limiting our ability to assess temporal dynamics across lactation. Second, dietary intake was measured using food frequency questionnaires, which are subject to recall bias and differences in measurement error across diet features. Third, the sample size for genetic analyses was modest relative to contemporary GWAS, limiting statistical power to detect weaker effects and rare variant associations. Finally, the cohort is comprised of U.S. mothers of predominately European ancestry with uncomplicated pregnancies who successfully breastfed for several months. Future studies in more diverse populations will be important for assessing the generalizability of these findings.

In summary, our study demonstrates that the human milk metabolome integrates information from maternal diet, genetics, and mammary gland physiology. By combining metabolomics with transcriptomic, genomic, and dietary profiling, we identify both environmental and genetic determinants of milk metabolites. Our findings demonstrate that maternal diet is the dominant measured determinant of variation across the human milk metabolome, while genetic variation reveals specific pathways governing mammary metabolism. These results establish a resource for future mechanistic studies of human lactation and provide a foundation for investigating how maternal nutrition and genetics shape the molecular composition of human milk and, ultimately, infant health.

## Materials and Methods

### Study cohort

This study made use of samples and data from the Mothers and Infants LinKed for Health Growth (MILK study), based at the University of Minnesota and University of Oklahoma Health Sciences Center. The study procedures have been extensively described in previous publications^26,64–66^. Briefly, participants who intended to exclusively breastfeed were enrolled prenatally during healthy, uncomplicated pregnancies at the University of Minnesota in collaboration with HealthPartners Institute (Minneapolis, MN) or the University of Oklahoma Health Sciences Center. Recruited mothers were 21–45 years old, non-smokers, and delivered singleton infants at full term (37 0/7 – 41 6/7 weeks gestation) with birth weight ≥2.5 kg and ≤4.5 kg. Clinical data for each mother-infant dyad were collected from their electronic health record and from electronic questionnaires at study visits at 1 and 6 months postpartum. The Institutional Review Boards of the University of Oklahoma, the University of Minnesota, and the HealthPartners Institute approved this study (STUDY00009021). This study has been registered with ClinicalTrials.gov (identifier NCT03301753). In the current study, all participants with the data type(s) required for each analysis were included.

### Milk sample collection

Milk samples were collected at study visits at approximately 1 month postpartum. Upon study visit arrival, participants fed their infants ad libitum from one or both breasts until infants were satisfied. Two hours following this feeding, milk was collected from the right breast using a hospital-grade electric breast pump (Medela Symphony; Medela, Inc., Zug, Switzerland), with expression ceasing when milk stopped flowing. Expressed milk volume and weight were recorded, milk was gently mixed, aliquots were made, and then stored at -80°C within 20 minutes of collection and kept at -80°C until thawed for compositional analyses.

### Milk macronutrient, cytokine, and hormone concentrations

Milk fat was separated from the aqueous phase by centrifugation, and skim milk was assayed using commercially available immunoassay kits for insulin, glucose, leptin, CRP, and IL6 and batch corrected as previously described^19,64–66^. Milk total fat, protein and carbohydrate concentrations were assessed by mid-infrared spectroscopy in a Miris Analyzer (Miris AB).

### Human milk metabolomics

The protocol to generate the milk metabolomics dataset used here has been extensively described^27,67,68^. 200 µL aliquots of 1-month milk samples were thawed on ice, vortexed, and incubated in extraction solvent at a 1:2 sample:solvent volume ratio −20 °C for 30 minutes. The extraction solvent was a mixture of isopropanol, acetonitrile, and water at a ratio of 3:3:2. Metabolites were processed by three analysis techniques: gas chromatography combined with high-resolution TOF MS (GC-MS), reversed-phase liquid chromatography coupled with high-resolution MS (LC-MS), and hydrophilic interaction chromatography with tandem mass spectrometry (LC-MS/MS). A quality control sample containing a standard mixture of amino and organic acids purchased from Sigma-Aldrich as certified reference material, was injected daily to perform analytical system suitability test and monitor recorded signals for day-to-day reproducibility. Samples were processed in 10 batches of 35 samples each, with 10 pooled milk samples and 40 external standards included to assess batch-to-batch variability.

Metabolites were annotated using known accurate masses and retention times and verified using in-house authentic standards. Missing values were imputed by replacement with 1/5 the limit of detection (the minimum recorded value for each metabolite). Data were combined from the three techniques into one data frame, with metabolites measured across more than one technique filtered by priority LC-MS/MS > LC-MS > GC-MS. 475 metabolites were identified, and metabolites with more than 20% missing values were removed from analyses, leaving 458 quantified metabolites.

Metabolite abundances were batch-corrected with ComBat^69^, log transformed, median-centered, and scaled to mean zero, standard deviation one. For all downstream analyses of metabolite values, we extracted the residuals of the normalized metabolite abundances after regressing on study site (OK vs. MN), infant age in days at sample collection, length of storage time between sample collection and metabolomics analysis, and volume of milk collected at the study visit. For analyses including genetic data, we additionally included the first three genetic principal components (PCs) when calculating the metabolite residuals. Metabolite subclass was assigned by lookup of the Human Metabolome Database ID (https://www.hmdb.ca/) provided by BPGbio for each metabolite (**Supplementary Table 1**).

### Dietary questionnaire

Maternal dietary intake data were collected via food frequency questionnaire. Earlier study participants used the Diet History Questionnaire II (DHQ II) during the third trimester of pregnancy and at one and three months postpartum; later study participants used the updated DHQ III once at one month postpartum. DHQ questionnaire responses were analyzed using Diet Calc (National Cancer Institute, Bethesda, MD, USA), and food and nutrient values were generated using the United States Department of Agriculture (USDA) Food Patterns Equivalent Database and the Food and Nutrient Database for Dietary Studies. Dietary feature values across DHQ II & III were integrated in the following manner: (1) filtered for the 211 dietary features quantified in both questionnaire versions; (2) filtered for features that had <90% zero values; (3) log normalized within each time point, with zeroes replaced with 1/10^th^ of the minimum non-zero value; (4) participant values averaged across time points, for those who took the DHQ II; (5) z-score standardizing feature values after grouping by each DHQ version and by the number of survey time points. Following this procedure there were no differences in mean value between survey version or the number of timepoints by participant.

To visualize the foods contributing to DHQ dietary features (e.g. **Figure 2B** right panel), we extracted food values from the DHQ III Nutrient Database^70^. For a given feature, we extracted the nutrient or food group value for females and portion size 1, and plotted the top 10 foods with value >0 (or fewer if less than 10 foods had non-zero values).

### Human milk RNA extraction, sequencing, and gene expression quantification

The human milk RNA extraction protocol, sequencing, and gene expression quantifications used in this study have been previously described^71^. RNA extraction, library preparation, and sequencing were performed at the University of Minnesota Genomics Center (UMGC). Briefly, bulk RNA was extracted from the whole milk cell pellet to profile gene expression of all cell types present in the milk sample. RNA was extracted from the cell pellet using the RNeasy Plus Universal HTP following the manufacturer’s instructions. RNA libraries were prepared with the TakaraBio Stranded Total RNA Pico Mammalian kit and sequenced on an Illumina NovaSeq 6000 S2 flow cell with 2×150 paired-end reads in two pools. Gene-level quantifications were generated using RNA-SeQC v2.3.4^72^.

### Maternal genotypes

This study utilized maternal genotypes derived from previously described whole-genome low-pass sequencing^19^. DNA was extracted from either human milk cells or saliva. Milk was collected at study visits and preserved as above. Saliva was collected in DNA Genotek Oragene 2 mL tubes following manufacturer’s instructions, either at study visits or at home and shipped to the lab. Milk and saliva specimens were shipped to Gencove for DNA extraction, library preparation, low-pass sequencing, and imputation and quality control in four batches. Genotype PCs were calculated by integrating our genotypes with the 1000 Genomes 30x coverage whole genome sequencing dataset^73^ as previously described^19^. We calculated pairwise relatedness using the “--make-rel” command in PLINK^74^ and iteratively removed one sample from pairs of participants with relatedness coefficient > 0.1.

### Statistical analyses

Statistical analyses were performed in RStudio^75^ (version 2024.12.0+467) unless otherwise noted. P-values were adjusted for multiple testing using the false discovery rate (FDR) method via the ‘qvalue’ R package^76,77^.

### Weighted gene co-expression network analysis

Weighted gene co-expression network analysis (WGCNA) was performed using the WGCNA package in R^78,79^. Starting with the normalized metabolite abundances adjusted for technical factors described above, hierarchical clustering of samples based on Euclidean distance was used to identify outlier samples. Two outlier samples were removed prior to network construction, leaving 347 samples and 458 metabolites for analysis. A signed weighted correlation network was constructed using the “blockwiseModules” function with a soft-thresholding power of 4, selected based on scale-free topology analysis using the “pickSoftThreshold” function. Modules were identified using dynamic tree cutting with a minimum module size of 10 metabolites and merged using a merge cut height of 0.25. Module eigengenes were calculated as the first PC of metabolite abundances within each module. Associations between module eigengenes and maternal clinical and milk traits were assessed using Pearson correlation. Enrichment of metabolite subclasses within each module was assessed by Fisher’s exact test.

### Lasso regression to identify maternal features influencing the milk metabolome

Lasso regression was performed for each milk metabolite as the outcome with three different sets of features as predictors: (1) dietary features derived from the DHQ data; (2) milk genes in the milk bulk transcriptomes; and (3) dietary features, milk metabolite QTL lead variants, and additional maternal metadata. Lasso regression was performed utilizing the glmnet R package^80^ with 10-fold cross validation. Ten replicate regressions were performed with different seeds, and features and weights from the replicate with highest correlation between out-of-fold predictions and observed metabolite abundances were selected for visualizations. Due to our limited sample size and goal of identifying the most strongly correlated features rather than predicting milk composition, we report the model R^2^ from the out-of-fold predictions rather than a held out test set. This can be interpreted as an optimistic estimate of predictive performance, because the same cross-validation procedure was used for model selection. Feature partial R^2^ to quantify the variance explained by individual features was calculated from a multiple regression model with features selected across >50% of replicates using the ‘rsq.partial’ command from R package ‘rsq’^81^. Feature p-values are the coefficient p-values from the multiple regression model. Details for the analysis of each feature set are described below.

(1) *Dietary features*: There were n=325 participants with both maternal diet data and milk metabolomics included in this analysis. Prior to lasso regression, dietary features were pruned by iteratively removing one feature from pairs with Pearson correlation > 0.99, retaining 174 features. Diet features and metabolite values were then rank normalized.
(2) *Milk gene expression*: There were n=299 participants with both 1-month milk transcriptome and metabolome data included in this analysis. Genes were filtered for protein-coding genes and those with TPM>0.1 in at least 25% of samples, retaining 13,795 genes. TPM values were standard normalized and then residualized on 13 technical covariates: 3’ bias, chimeric alignment rate, high quality ambiguous alignment rate, median of transcript coverage, sample RIN, RNA concentration, rRNA reads, total reads, RNAlater preservation status, unique vendor QC passed reads, and sequencing batch.
(3) *Combined feature set (diet, metabolite QTL, and maternal clinical features)*: We combined DHQ features described in (1) above with maternal genotypes for the lead variant at each of 35 genome-wide significant metabolite QTL, and the following maternal features: maternal age at recruitment, gestational diabetes status, maternal pre-pregnancy BMI, cesarean vs. vaginal delivery, and parity for n=183 unrelated participants with data for all features and milk metabolomics. We regressed 3 genetic PCs and technical covariates (study site, infant age in days, milk storage time, milk volume expressed) from log normalized metabolite abundances prior to lasso regression.

### GWAS of milk metabolites

There were n=197 unrelated participants with 1-month milk metabolomics and genotype data included in each metabolite GWAS. To prepare the metabolomics data, we log normalized metabolite abundances and then regressed out the first 3 genetic PCs from the joint PCA with 1000 Genomes individuals and the first 10 PCs of the log normalized metabolite matrix in a linear regression. We regressed out metabolite PCs to remove the effects of hidden confounders and improve our power to detect genetic effects, as is common in molecular QTL mapping^82^. We then performed GWAS via linear mixed model in GEMMA v0.98.5^83^. We considered genetic associations with a stringent minor allele frequency of >10% due to our limited sample size. We identified genome-wide (p<5×10^-8^) and study-wide (p<1.09×10^-10^, correcting for 458 metabolites) and significant loci by grouping variants within 1 Mb that fell below the p-value threshold.

To compare genetic effects on milk metabolites to plasma metabolites, we downloaded study-wide significant associations reported in the supplementary tables of plasma metabolite GWAS from the Canadian Longitudinal Study on Aging^37^ and the EPIC-Norfolk study^49^. We identified shared metabolites between those studies and our dataset by matching of HMDB, KEGG, and/or PubChem IDs. We then extracted the milk genetic effects from our GWAS summary statistics for the same metabolite reported in the plasma study. Power to replicate plasma QTL associations in milk was estimated under an additive genetic model, assuming that the true per-allele effect in milk was equal to the effect reported in plasma. For each association, we calculated the expected noncentrality parameter of the Z statistic as

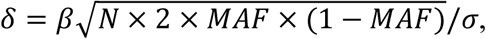

where *β* is the reported plasma effect size, *N* is the milk sample size, *MAF* is the minor allele frequency in the MILK cohort, and *σ* is the metabolite-specific residual standard deviation in the MILK cohort. Power was calculated as *P*(| *Z* |> *z*1 − *a*/2) for *Z* ∼ *N*(*δ*, 1), using a two-sided significance threshold of *a* = 0.05.

To assess the predictive ability of plasma metabolite PRS on milk metabolites in our sample, we downloaded PGS score files for plasma metabolites (quantified by Metabolon) from OmicsPred^41^. We calculated PGS for our n=197 unrelated participants with genotype and milk metabolomics data for 135 metabolites identified as present in both our dataset and the OmicsPred data by manual matching. PGS were calculated with the OmicsPred score file via the ’--score’ command in PLINK 2.0^84^. We calculated the variance explained in each milk metabolite by its plasma PGS by regressing the input phenotype values for the metabolite GWAS (i.e. log normalized abundances with 3 genetic PCs and 10 metabolite PCs regressed out) against PGS. Not all genetic variants in the PGS were available in our dataset, and we observed a significant correlation between the proportion of available SNPs and the ratio of milk to plasma R^2^ (Spearman’s rho = 0.24, p-value = 0.0058). We accounted for this in our estimate of power to replicate the PGS signal in milk by scaling the internal PGS R^2^ in plasma by the proportion of PGS variants available in our dataset, and using this as the expected R^2^ in milk. We then calculated replication power as the power to reject the null hypothesis of 1 degree of freedom F-test with n=197, alpha=0.05 using the ‘f.test.power’ function of the *pwrss* package in R^85^.

To assess the effect of lead variants at putative milk-specific mQTL on the same metabolites in other tissues, we downloaded genome-wide summary statistics from the GWAS catalog^86^ with the following accession numbers: GCST90243465 (plasma xanthine)^44^, GCST90243463 (plasma hypoxanthine)^44^, GCST90200314 (plasma N-acetyl neuraminic acid)^37^, GCST90318854 (cerebrospinal fluid N-acetyl neuraminic acid)^47^.

### Mediation analysis with two-stage least squares regression

To test if gene expression mediated an association between a metabolite and genetic variant in our cohort (e.g. rs164745, *DPEP1* expression, and cysteine-glycine abundance), we performed two-stage least squares regression using the ‘ivreg’ package in R^87^. In this analysis we used the normalized and residualized metabolite abundances used in the metabolite GWAS above. We used gene expression values that were standard normalized and then residualized on 13 technical covariates (3’ bias, chimeric alignment rate, high quality ambiguous alignment rate, median of transcript coverage, sample RIN, RNA concentration, rRNA reads, total reads, RNAlater preservation status, unique vendor QC passed reads, and sequencing batch) and the first 3 genetic PCs.

### Colocalization of metabolite QTL and eQTL

Colocalization was performed between metabolite QTL summary statistics from this study, eQTL summary statistics from human milk^19^, and GTEx v8 eQTL summary statistics^43^ using the *coloc* R package^88,89^. Summary statistics were filtered to variants within 500kb of the metabolite QTL lead SNP. QTL signals were fine-mapped using the ‘runsusie’ command. LD matrices were generated using PLINK’s ‘--r square’ function^74^ with MILK study genotype data. Colocalization was run with the command ‘coloc.susie’^90^ with a prior probability of colocalization of p_12_=3.5×10^-5^. For traits where fine mapping identified more than one credible set at a locus, the trait pair with the highest value of PP.H4 was used for plotting (e.g. **Figure 5D**).

### Assessment of sialic acid QTL effects on HMOs in MILK and CHILD cohorts

We tested for associations between rs71876558 and HMO concentrations in the MILK cohort using the same GWAS pipeline as for milk metabolites described above. HMO concentrations in the MILK cohort were described previously^19,91^. Summary statistics from an HMO GWAS in the CHILD cohort study^13^ were downloaded from the GWAS catalog^86^ (accession numbers GCST90320263-GCST90320285). While the lead milk sialic acid GWAS variant (rs71876558) was not available in these summary statistics, we utilized summary stats for variant rs10972852 which is in strong LD (r^2^ = 0.978) with rs71876558 in Europeans^92^.

## Acknowledgements

We thank all MILK study staff and participants for their contributions, and members of the Albert and Blekhman labs for helpful discussions related to this project. This work was supported by the resources and staff at the University of Minnesota Genomics Center (https://genomics.umn.edu). This work was carried out in part by resources provided by the Minnesota Supercomputing Institute (https://www.msi.umn.edu/).

## Author contributions

Conceptualization: KEJ, FWA, RB, EWD

Methodology: KEJ, JJAH, MAK, AJ, EMN, FWA, RB, EWD

Investigation: KEJ, JJAH, MAK, KP, SP, SW

Formal analysis: KEJ, RD, AY

Funding acquisition: KEJ, LB, EFL, EMI, DAF, FWA, RB, EWD

Supervision: KEJ, KP, FWA, EWD, RB

Writing - original draft: KEJ

Writing - review and editing: KEJ, RD, AY, JJAH, AJ, MAK, EMN, KP, SP, SW, LB, EFL, EMI, DAF, FWA, RB, EWD

## Data availability

Code and data to reproduce the analyses and figures are deposited at https://github.com/kelsj/human_milk_metabolomics. RNA and DNA sequencing data utilized in this paper are deposited at dbGaP (Study Accession: phs003408.v1.p1). Milk metabolite GWAS summary statistics are available in the NHGRI-EBI GWAS catalog (accessions GCST90999369 - GCST90999826).

