## supplemental_figures for "Maternal diet and genetics shape the human milk metabolome"

#### Supplementary Figure Legends

**Supplementary Figure 1. Quality control of human milk metabolomics. A)** Boxplots of coefficient of variation (CV) between replicate pooled QC samples (light green) and test samples (dark green). **B)** Scatter plot of mean metabolite abundance (x-axis) and CV in QC samples. Metabolites colored by subclass.

**Supplementary Figure 2. Correlations between WGCNA module 3 and milk composition. A)** Scatter plot of milk sample WGCNA module 3 eigengenes (x-axis) and IL-6 protein concentration (y-axis). **B)** Scatter plot of milk sample WGCNA module 3 eigengenes (x-axis) and glucose concentration (y-axis).

**Supplementary Figure 3. Correlations of milk metabolite module 1 acylcarnitines with milk IL-6 and glucose concentrations.** Each dot represents the Pearson correlation coefficient for one of 23 acylcarnitines in metabolite module 1 with milk concentrations of IL-6 protein (right) or glucose (left). Dots are colored by the fatty acyl chain length: short-chain (C2–C5), medium-chain (C6–C12), and long-chain (≥C14). Dot size represents the -log10(p-value) of the correlation coefficient.

**Supplementary Figure 4. Pairwise correlations between diet features and milk metabolites by metabolite subclass.** Each bar represents the number of significant pairwise correlations (q-value < 0.05) between a milk metabolite and a dietary feature. Each bar is a metabolite subclass, and color correspond to subclass. Bars to the left signify negative correlations, and bars to the right signifiy positive correlations.

**Supplementary Figure 5. Milk metabolites correlated with HEI total score.** Scatter plots showing the relationship between milk N-acetylornithine (left panel) or riboflavin (right panel) with Healthy Eating Index (HEI) total score. Higher HEI total score indicates better adherence to dietary guidelines.

**Supplementary Figure 6. Variation explained in HEI scores by the milk metabolome.** Variation explained in each HEI component (out-of-fold $R^2$) by the milk metabolites selected by lasso regression model. Each vertical bar is an HEI component, and bar color represents the number of metabolites selected by the model. The dotted horizontal line represents the median $R^2$.

**Supplementary Figure 7. Colocalization of *DPEP1* eQTL and cysteine-glycine mQTL.** Scatter plot of *DPEP1* eQTL p-values (x-axis) and cysteine-glycine QTL (y-axis) in milk.

**Supplementary Figure 8. Plasma metabolite genetic associations that did not replicate in milk.** Forest plot displaying genetic effect estimates and 95% confidence intervals for metabolite QTL in plasma (dark purple) and milk (teal). The table lists the likely causal gene, lead variant in plasma, and metabolite. Associations were included here if they had >90% power to replicate the effect seen in plasma (from Chen et al., 2023) and p-value > 0.2, in milk.

**Supplementary Figure 9. Effect estimates of plasma metabolite PGS for the same metabolite in milk.** Forest plot displaying the effect estimates and 95% confidence intervals of plasma PGS on milk metabolite abundance. Point estimates and error bars are colored by the PGS $R^2$ in plasma, and ordered by the effect estimate in milk.

**Supplementary Figure 10. N-acetylornithine abundance in milk vs. plasma PGS.** Scatter plot of N-acetylornithine normalized abundance (y-axis) vs. PGS for N-acetylornithine in plasma.

**Supplementary Figure 11. Genetic associations with milk metabolites at the *PDE6A* locus.** Forest plot displaying the estimated effect estimate and 95% confidence interval for each milk metabolite with a nominal, genome-wide significant, or study-wide significant association at rs17725010.

**Supplementary Figure 12. Fine-mapping of milk metabolite associations at the *PDE6A* locus.** Posterior inclusion probabilities (PIP) after fine-mapping the six milk metabolites with genome-wide or study-wide significant genetic associations at the *PDE6A* locus. Lollipops represent PIPs for each metabolite at each variant, colored by milk metabolite. Variants are ordered by their position along chromosome 5 (hg38).

**Supplementary Figure 13. Distribution of *PDE6A* expression across GTEx tissues and human milk.** TPM values from GTEx and human milk cells are displayed as box plots.

**Supplementary Figure 14. eQTL effects across tissues of the lead variant at the *PDE6A* milk mQTL locus.**

**Supplementary Figure 15. Pairwise correlations between abundances of purine metabolism pathway metabolites in milk.**

**Supplementary Figure 16. Colocalization of milk metabolite QTL and GTEx eQTL at the *PDE6A* locus.**

**Supplementary Figure 17. Distribution of *GNE* expression across GTEx tissues and human milk.** TPM values from GTEx and human milk cells are displayed as box plots.

**Supplementary Figure 18. Neu5Ac abundance in milk vs. plasma PGS.** Scatter plot of Neu5Ac normalized abundance (y-axis) vs. PGS for Neu5Ac in plasma.

**Supplementary Figure 19. Associations between a variant tagging the Neu5Ac QTL at the *GNE* locus with HMOs in the MILK and CHILD cohort studies.** Forest plot displaying the genetic effect estimates and 95% confidence intervals for 22 HMO traits in the MILK (left) and CHILD (right) cohort studies. The lead variant for Neu5Ac at the *GNE* locus (rs71876558) was not available in CHILD cohort summary statistics, so we selected rs10972852, which is in strong LD with rs71876558 ($r^2$ = 0.978 in Europeans).

**Supplementary Figure 20. Correlation between TMAO in milk and its top lasso-selected dietary feature.** Left panel: scatter plot displaying the normalized dietary intake of PFA 22:5 (docosapentaenoic acid, DPA) (x-axis) vs. milk TMAO (trimethylamine N-oxide). Right panel: top foods contributing to dietary docosapentaenoic acid (22:5)**.** Each bar represents the listed food's PFA 22:5 concentration, according to the DHQ index.

**Supplementary Figure 1. Quality control of human milk metabolomics.**

A.

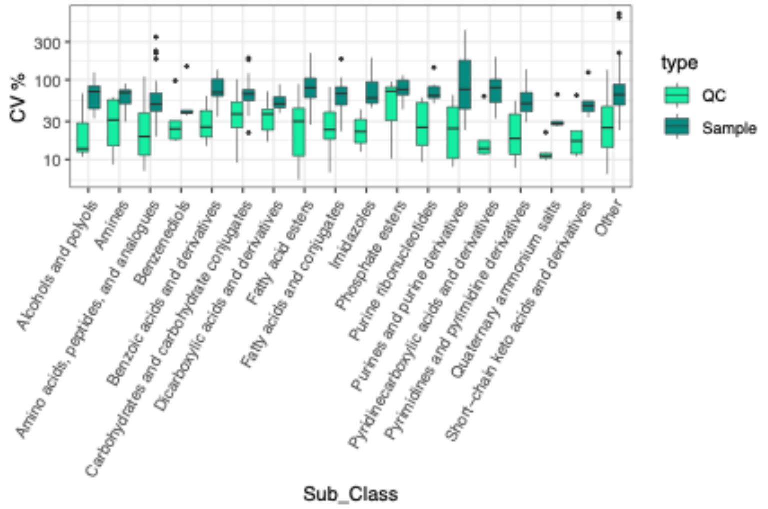


B.

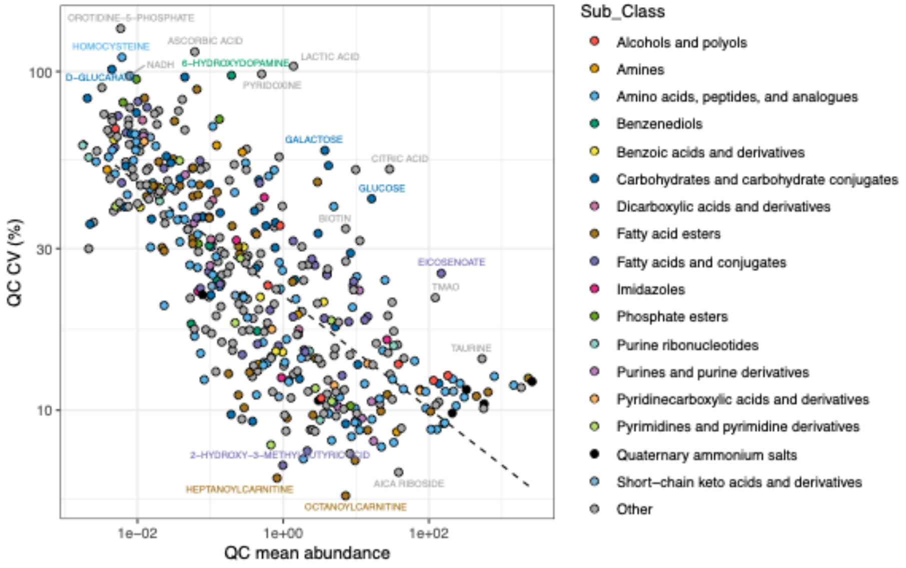


**Supplementary Figure 2. Correlations between WGCNA module 3 and milk composition.**

A.

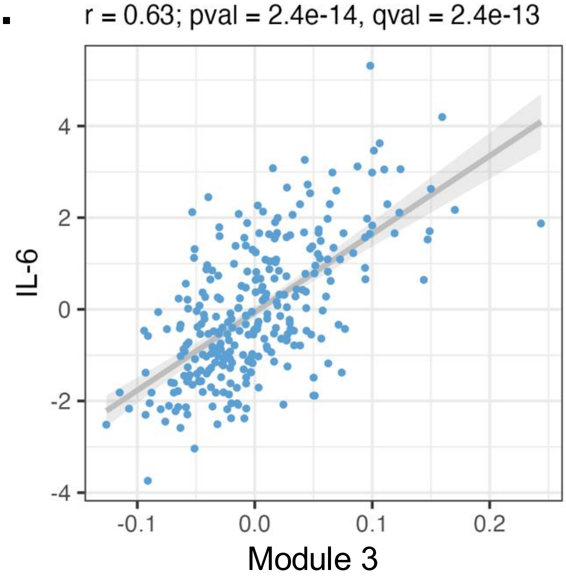


B.

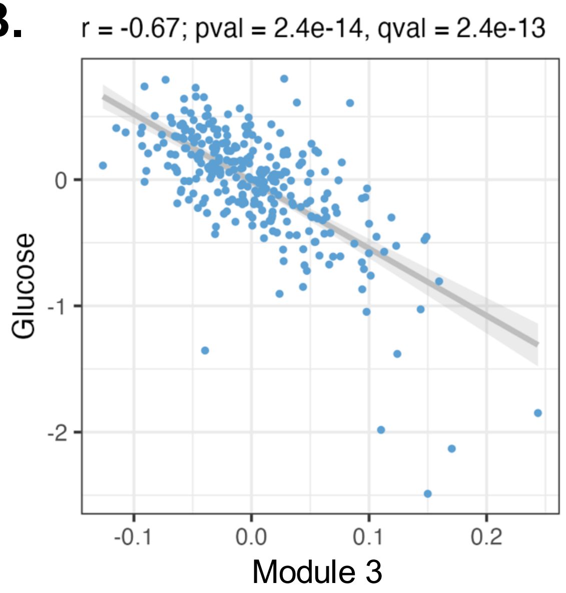

**Supplementary Figure 3. Correlations of milk metabolite module 1 acylcarnitines with milk IL-6 and glucose concentrations.**

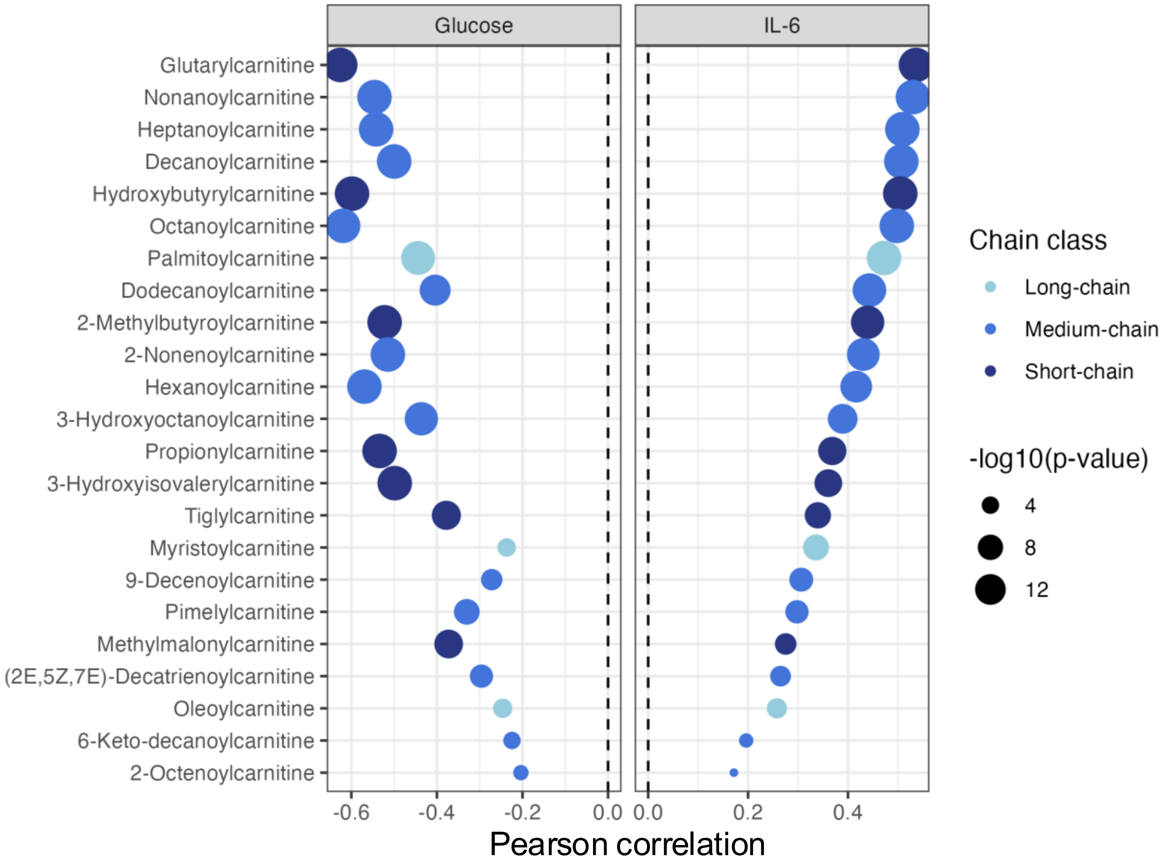

**Supplementary Figure 4. Pairwise correlations between diet features and milk metabolites by metabolite subclass.**

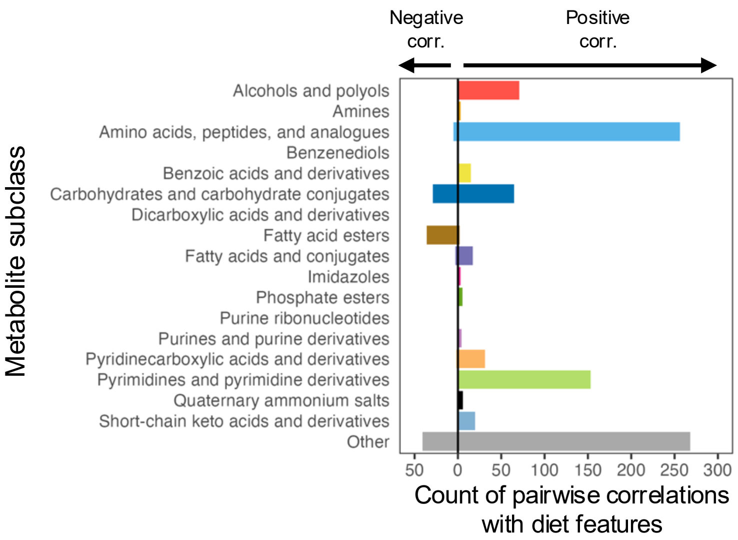


**Supplementary Figure 5. Milk metabolites correlated with HEI total score.**

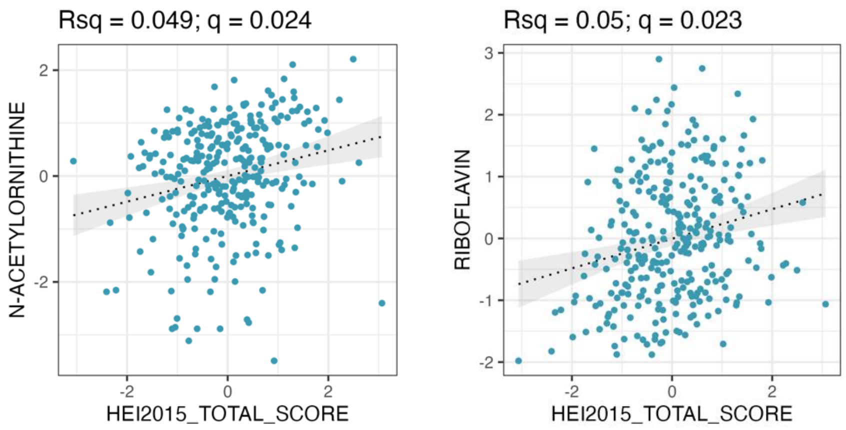

**Supplementary Figure 6. Variation explained in HEI scores by the milk metabolome.**

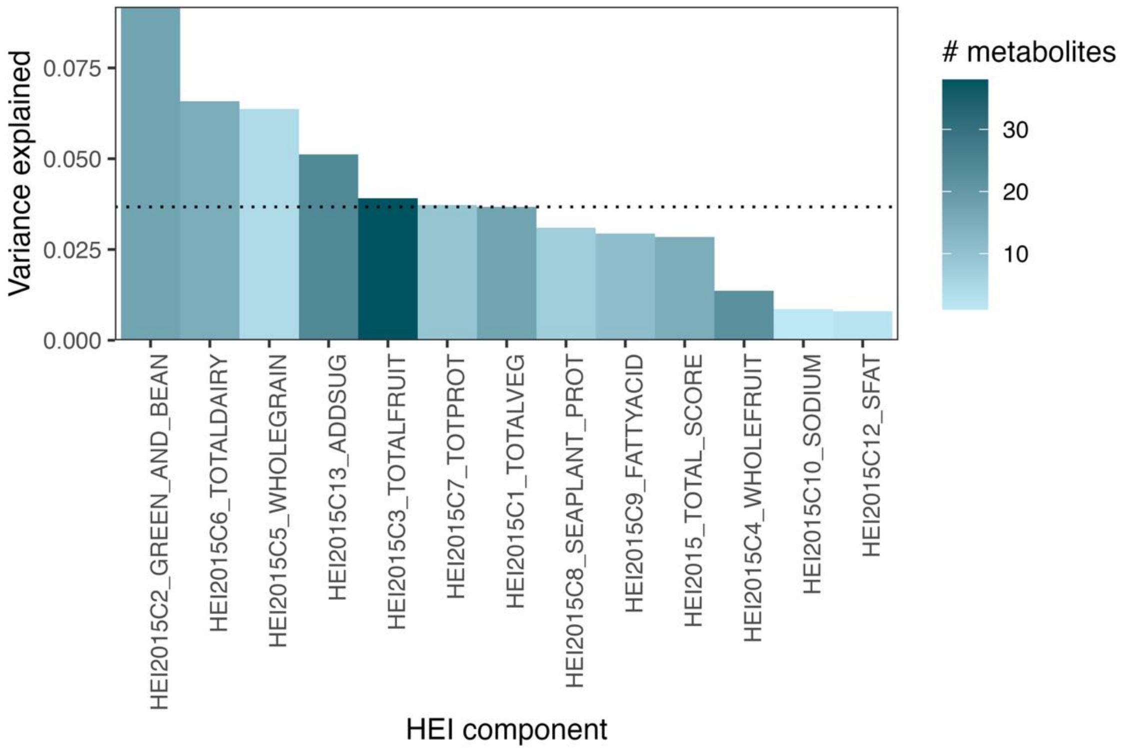

**Supplementary Figure 7. Colocalization of *DPEP1* eQTL and cysteine-glycine mQTL.**

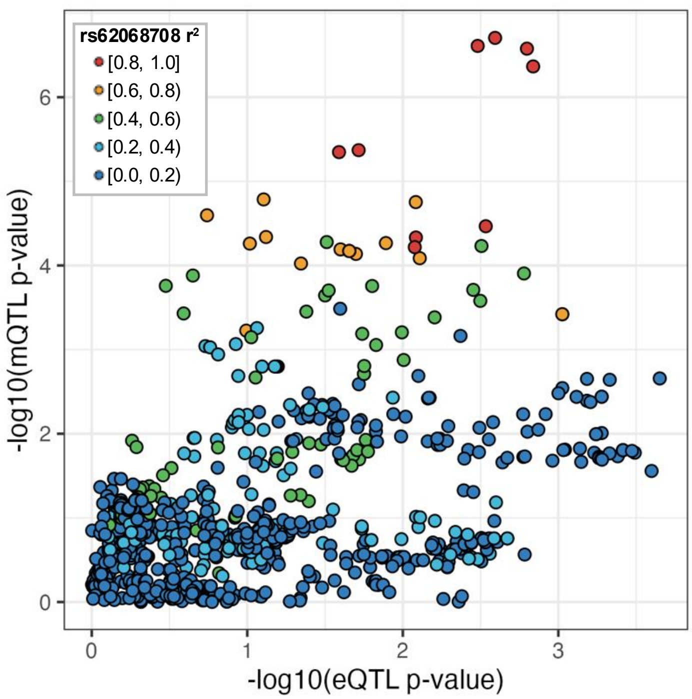

#### Supplementary Figure 8. Plasma metabolite genetic associations that did not replicate in milk.

| Causal gene | Lead variant | Metabolite |
|---|---|---|
| *NAT2* | 8_18414867_T_C | N-acetyl putrescine |
| *AGXT2* | 5_35045721_G_A | 3-amino isobutanoic acid |
| *NAT8* | 2_73650092_G_A | N-acetyl glutamine |
| | | N-acetyl asparagine |
| | | N-acetyl-L-tyrosine |
| | | 2-Aminooctanoic acid |
| *UGT1A3* | 2_233755940_ATC_A | Bilirubin |
| *ACADM* | 1_75758325_C_T | Nonanoylcarnitine |
| | | 9-Decenoylcarnitine |
| *SLCO1B1* | 12_21178615_T_C | Tetradecanedioate |
| | | Hexadecanedioate |
| *ACADS* | 12_120738280_G_A | Methyl succinic acid |
| | | Ethylmalonic acid |

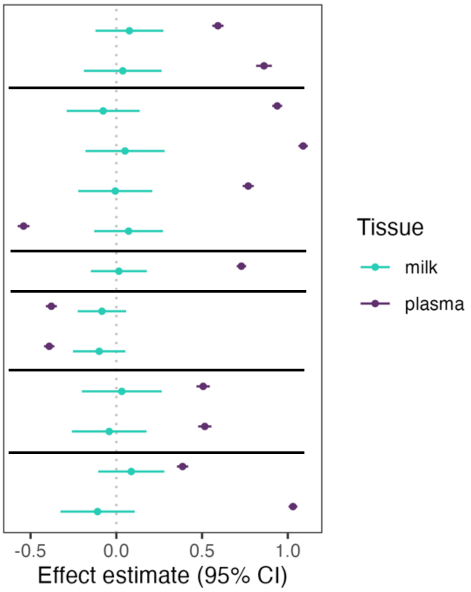


#### Supplementary Figure 9. Effect estimates of plasma metabolite PGS for the same metabolite in milk.

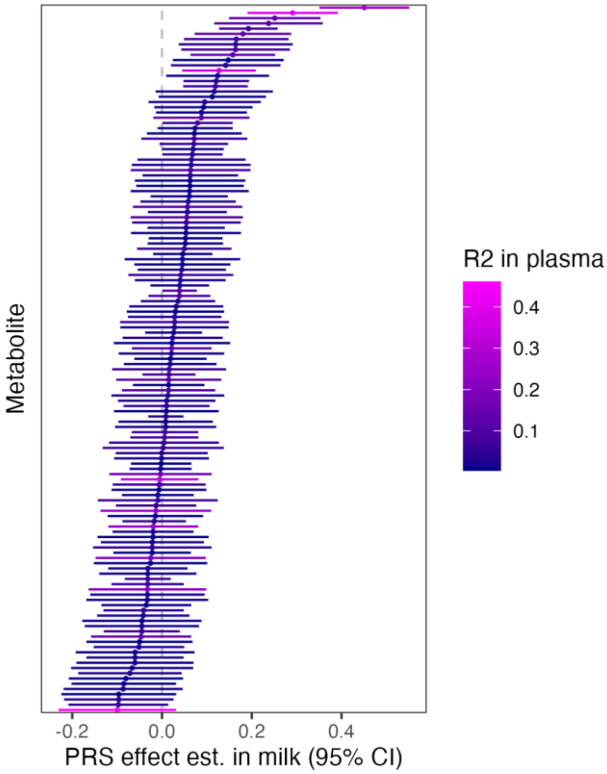

### Supplementary Figure 10. N-acetylornithine abundance in milk vs. plasma PGS.

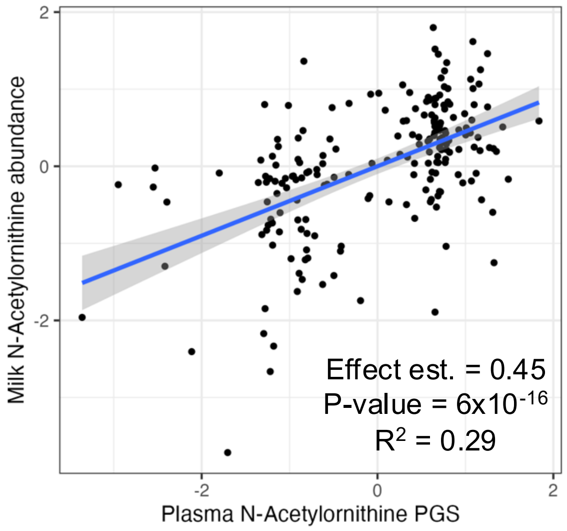

#### Supplementary Figure 11. Genetic associations with milk metabolites at the *PDE6A* locus.

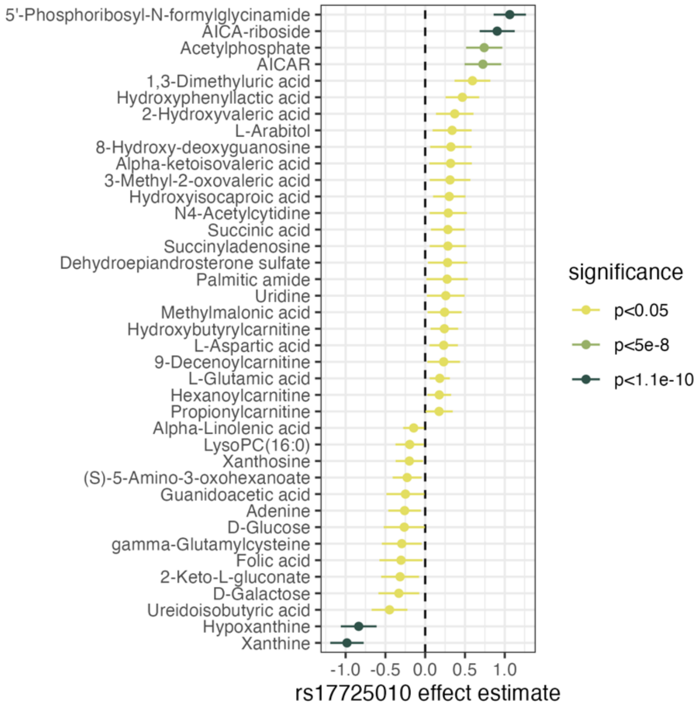


#### Supplementary Figure 12. Fine-mapping of milk metabolite associations at the *PDE6A* locus.

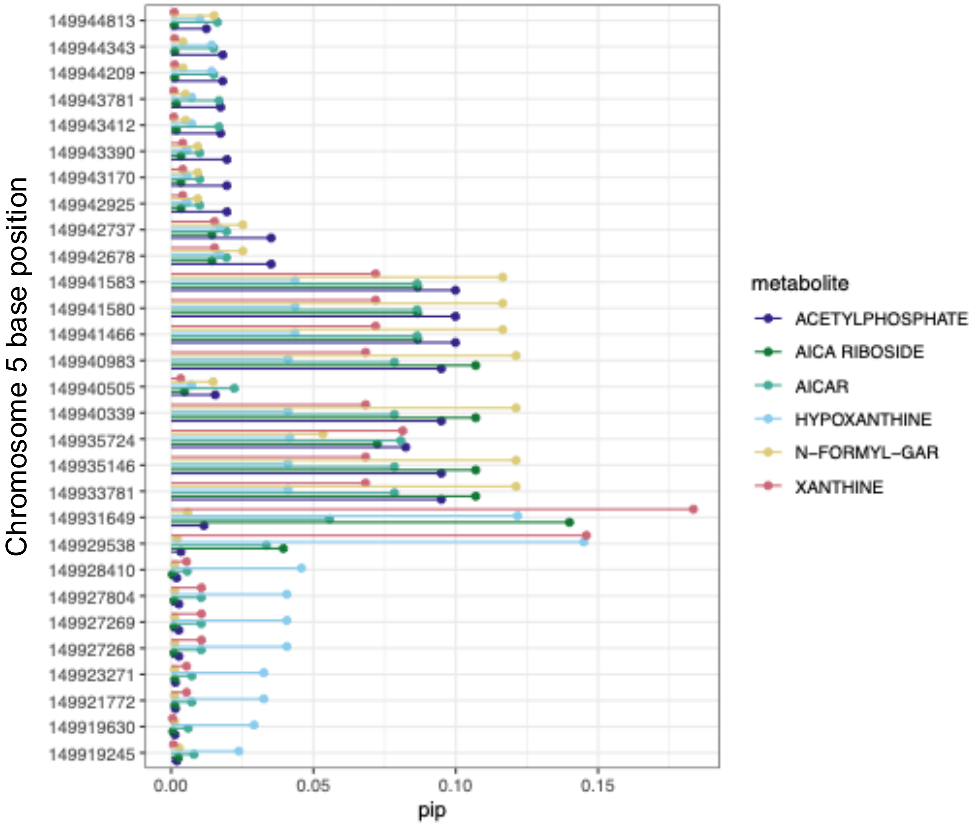

#### Supplementary Figure 13. Distribution of *PDE6A* expression across GTEx tissues and human milk.

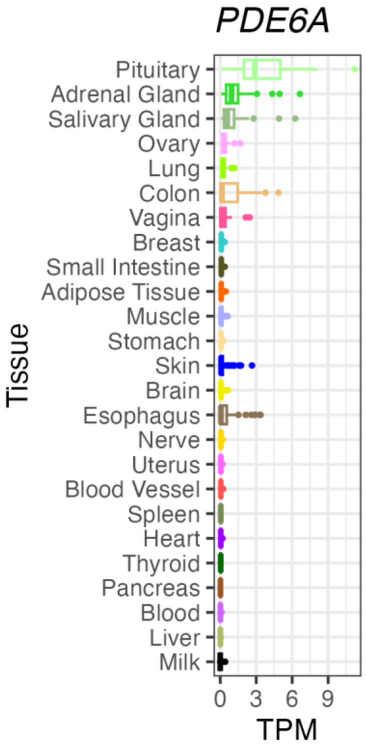


#### Supplementary Figure 14. eQTL effects across tissues of the lead variant at the *PDE6A* milk mQTL locus.

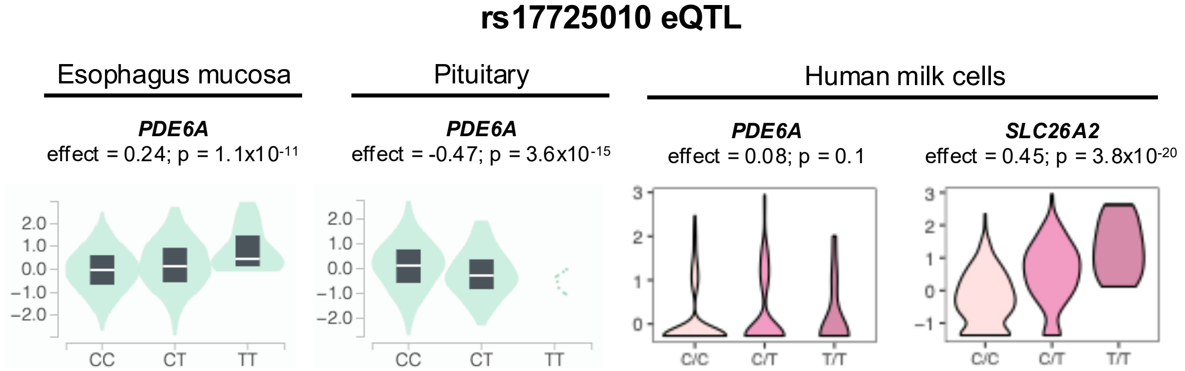

#### Supplementary Figure 15. Pairwise correlations between abundances of purine metabolism pathway metabolites in milk.

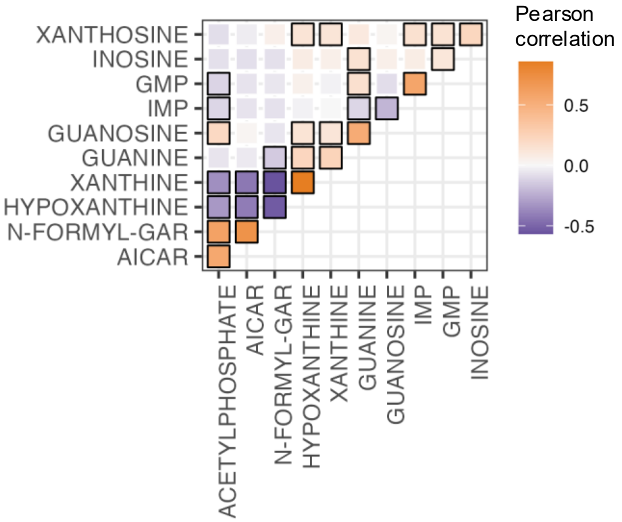


#### Supplementary Figure 16. Colocalization of milk metabolite QTL and GTEx eQTL at the *PDE6A* locus.

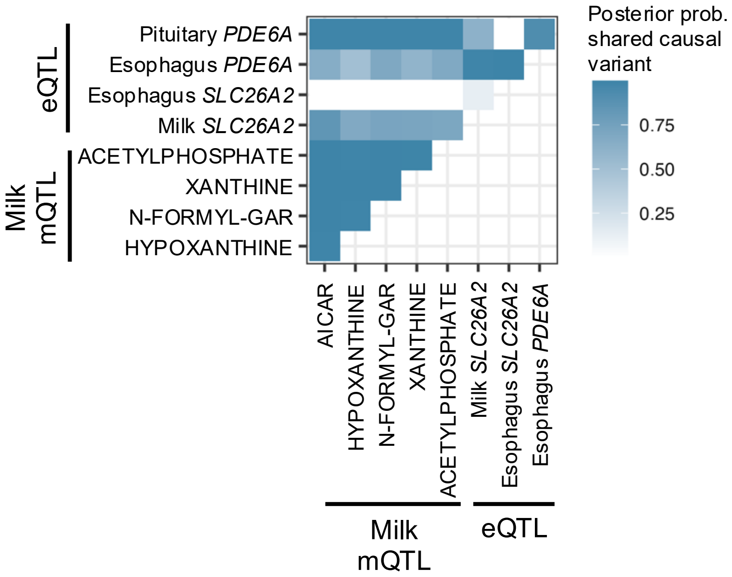

**Supplementary Figure 17. Distribution of *GNE* expression across GTEx tissues and human milk.**

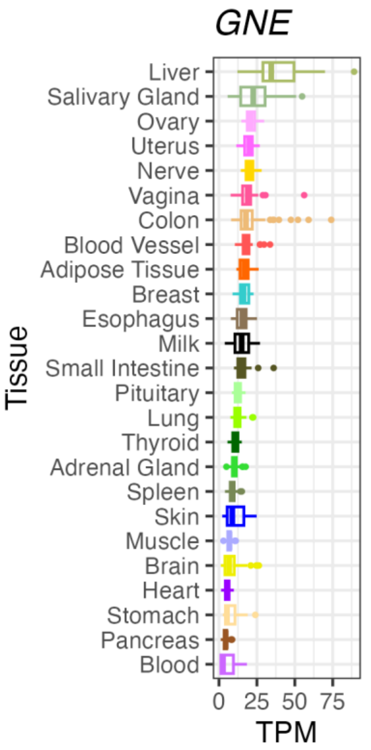


**Supplementary Figure 18. Neu5Ac abundance in milk vs. plasma PGS.**

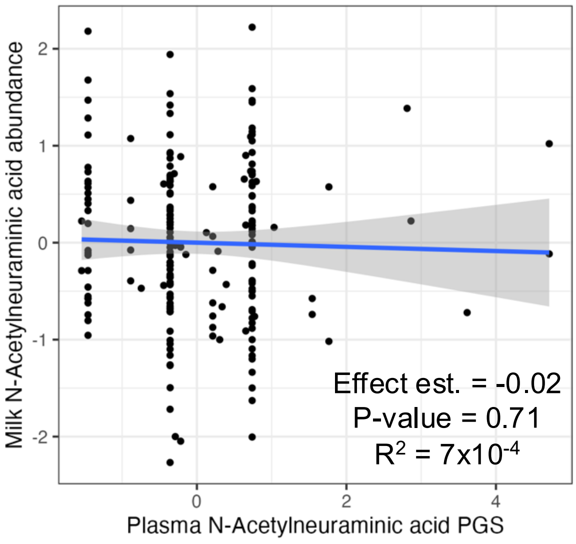

**Supplementary Figure 19. Associations between a variant tagging the Neu5Ac QTL at the *GNE* locus with HMOs in the MILK and CHILD cohort studies.**

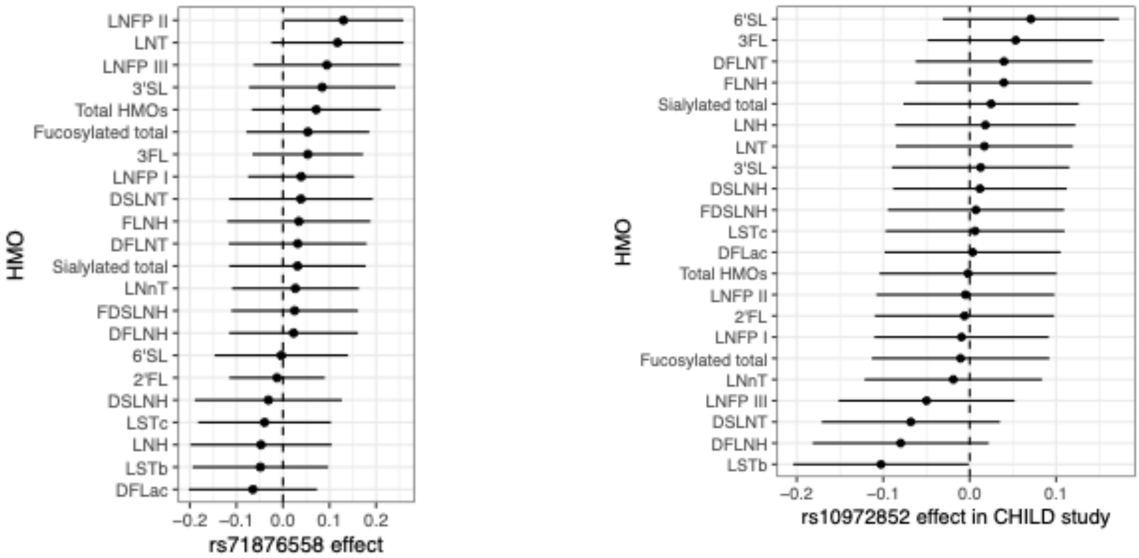

#### Supplementary Figure 20. Correlation between TMAO in milk and its top lasso-selected dietary feature.

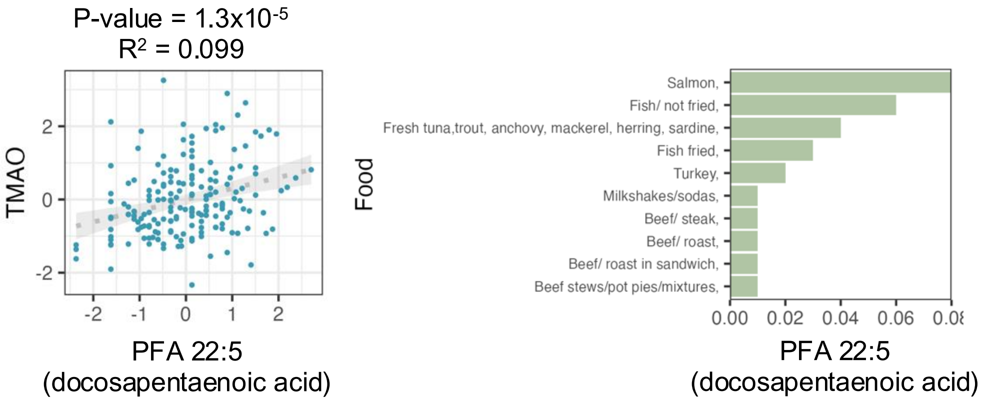
